# RNA interactions with CTCF are essential for its proper function

**DOI:** 10.1101/530014

**Authors:** Ricardo Saldaña-Meyer, Javier Rodriguez-Hernaez, Mayilaadumveettil Nishana, Karina Jácome-López, Elphege P. Nora, Benoit G. Bruneau, Mayra Furlan-Magaril, Jane Skok, Danny Reinberg

## Abstract

The function of the CCCTC-binding factor (CTCF) in the organization of the genome has become an important area of investigation, but the mechanisms of how CTCF dynamically contributes to genome organization is not clear. We previously discovered that CTCF binds to large numbers of endogenous RNAs; promoting its oligomerization. Here we found that inhibition of transcription or interfering with CTCF ability to bind RNA through mutations of two of its 11 zinc fingers that are not involved with CTCF binding to its cognate site in vitro, zinc finger-1 (ZF1) or −10 (ZF10), disrupt CTCF association to chromatin. These mutations alter gene expression profiles as CTCF mutants lose their ability to promote local insulation. Our results highlight the importance of RNA as a structural component of the genome, in part by affecting the association of CTCF with chromatin and likely its interaction with other factors.

**Bullet Points (4 of max 75 words):**

- Transcriptional inhibition disrupts CTCF binding to chromatin
- RNA-binding regions (RBR) in CTCF are found within ZF1 and ZF10
- Local insulation is markedly decreased in ZF1∆ and ZF10∆ mutant rescues
- Gene expression and chromatin organization are disrupted by RBR mutants

## Introduction

Spatial and temporal control of gene expression is crucial for the development of multicellular organisms. Improper gene expression leads to developmental abnormalities and diseases such as cancer. In addition to the “linear” genetic information, the three dimensional (3D) spatial organization of the eukaryotic genome within the nucleus contributes to genome function (Bonev and Cavalli, 2016; Merkenschlager and Nora, 2016).

The 3D genome is hierarchically organized: from nuclear compartments, to large topologically insulated domains (TADs), to short-range cis-interactions. TADs are formed primarily by the interaction of CTCF, with the cohesin complex, and likely other proteins (Havva et al., unpublished results). CTCFand the cohesin complex are pivotal to 3D structure formation (Rowley and Corces, 2018). The depletion of either factor has drastic effects on chromatin structure wherein TADs essentially disappear (Nora et al., 2017; Rao et al., 2017). The most widely accepted explanation of how chromatin organizes 3D structure is through the loop extrusion model (Fudenberg et al., 2017). This model proposes that cohesin rings create loops by actively extruding DNA until the cohesin complex finds two CTCF binding sites in convergent orientation. This explains the underlying mechanism of TAD organization, but many questions remain unanswered about how these domains are regulated temporal and in a cell-type specific manner. Equally important is that within these (“mega”) TADs there are intra (“mini”) TADs, which are also insulated chromatin domains. These intra-TADs are insulated, as described above (“mega” TADS), but also require the interaction of transcription factors.

Although almost all TAD boundaries are enriched by CTCF and cohesin, the majority of CTCF-bound sites are found elsewhere in the genome (Merkenschlager and Nora, 2016). Furthermore, CTCF and cohesin binding sites are significantly conserved among cell types, but still many of them, as well as many TADs, display cell-type specific patterns and changes during differentiation as a result of stage-specific transcription factors (Pekowska et al., 2018; Stadhouders et al., 2018).

Together with CTCF, YY1, cohesin and Mediator complexes are implicated in defining chromatin architecture at different topological ranges and all of these proteins bind RNA (Lai et al., 2013; Li et al., 2013; Phillips-Cremins et al., 2013; Saldaña-Meyer et al., 2014; Sigova et al., 2015). However, a growing number of examples demonstrate that RNA can recruit and stabilize or destabilize protein binding to chromatin, as in the case of the PRC2 complex and YY1, respectively (Beltran et al., 2016; Kaneko et al., 2014a; 2014b; Sigova et al., 2015). Furthermore, both CTCF and YY1 can form dimers and oligomers in an RNA-dependent manner which may account for the regulation of far cis-interactions on chromatin (Saldaña-Meyer et al., 2014; Weintraub et al., 2017).

Here, we sought to identify the functional relevance of CTCF-RNA interactions using two strategies: 1) inhibiting transcription and 2) rescuing the loss of wild-type endogenous CTCF with RNA binding-deficient mutants. We concentrated on three distinct levels of regulation: 1) gene expression using single cell RNA-Seq and bulk RNA-Seq, 2) chromatin binding via ChIP-Seq and 3) chromatin structure via 5C and Hi-C. We demonstrate that the lack of RNA binding to CTCF disturbs its stability on chromatin with direct and indirect effects on gene expression and 3D chromatin organization.

## Results

### Transcriptional Inhibition disrupts CTCF binding to chromatin

To unbiasedly test if RNA binding is integral to CTCF activity genome-wide, we first performed chromatin immunoprecipitation followed by deep-sequencing (ChIP-Seq) after transcriptional inhibition (TI) with co-incubation of DRB and Triptolide in mouse embryonic stem cells (mESCs). The treatment inhibits both elongation and initiation of transcription and promotes degradation of RNAP II (Bensaude, 2014), yet it revealed no influence on CTCF protein abundance (**Figure S1A)**. Nonetheless we detected an overall decrease in binding to chromatin genome-wide primarily at transcriptional start sites (TSSs) (**Figure 1A; Figure S1B-C)**. Since CTCF is widely present throughout the genome within both intragenic and intergenic regions, we next focused on the specific genomic distribution between individual CTCF binding sites in the control versus the sites whose binding was significantly decreased in TI. We found that CTCF binding sites within TSSs and promoters were the most significantly affected class (**Figure 1B-C: Figure S1D**). Remarkably, the sites perturbed by transcriptional inhibition were those with motifs with significantly lower affinity when compared to a random sample of stable CTCF bound regions (Mann-Whitney test, p < 0.0001; **Figure 1D**). Looking at chromatin structure, 5C experiments targeting a 4 MB region showed that the CTCF-CTCF loop at the *HoxA* cluster boundary was disrupted (**Figure 1E; top & middle**) without the loss of CTCF in the boundaries illustrated by the overlapped ChIP-seq tracks (**Figure 1E; bottom**). However, elsewhere in the region CTCF binding was decreased and we observed a general increase of chromatin interactions (**Figure S1E**). Similar results were observed after depleting RNA by incubation with RNase A (**Figure S1F**). These results favor the hypothesis that CTCF binding to chromatin as well as CTCF-CTCF interactions (Saldaña-Meyer et al., 2014) are stabilized by RNA molecules at defined sites within the genome.

**Figure 1.**
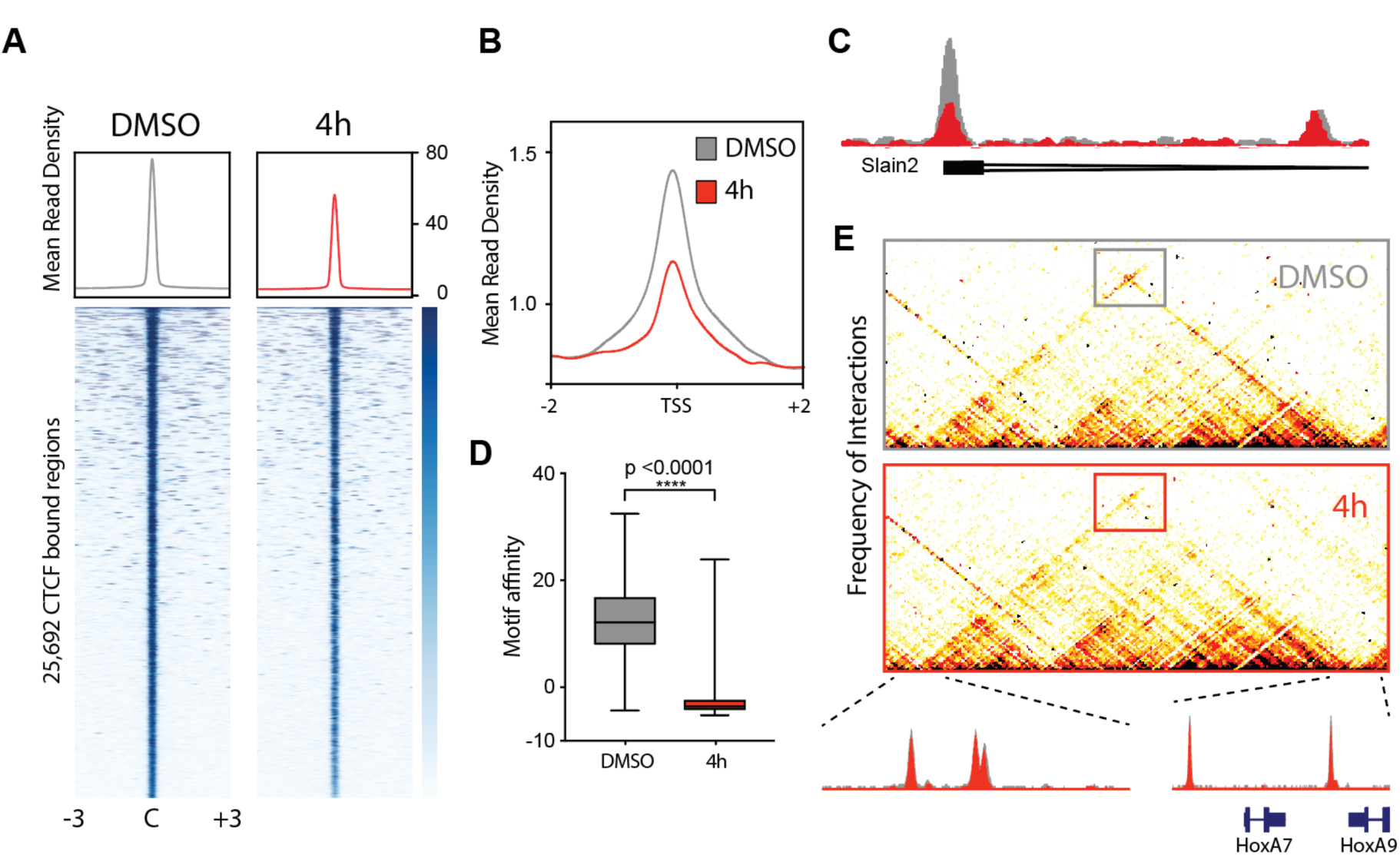
Transcriptional Inhibition disrupts CTCF binding predominantly in TSS. Transcription was inhibited in mESCs for 4 hours with co-incubation of DRB and Triptolide. Cells incubated with DMSO served as control. **(A)** Shows CTCF ChIP-seq heatmaps centered and rank-ordered on CTCF binding sites. Corresponding average density profiles are plotted at the top of the heatmaps to illustrate the differences between DMSO and 4 h of TI. **(B)** Average density profiles for the same ChIP-Seq as **A** but centered on TSS **(C)** Representative example of a CTCF peak with decreased binding to the TSS of the Slain2 gene. ChIP-Seq tracks for DMSO (grey) and 4 h of TI (red) are overlapped. **(D)** Box plot showing the motif affinity scores for CTCF binding sites lost after transcriptional inhibition versus a random set of CTCF binding sites in the control condition**. (E)** 5C heatmap depicting the interaction frequency between restriction fragments across a 1-Mb region surrounding the *HoxA* cluster. Darker colors represent increasing interaction frequency. The dotted lines zoom into overlapped ChIP-Seq tracks for DMSO (grey) and 4 h of TI (red) illustrating no change in CTCF binding for the loop enclosed in a rectangle.

### High-resolution mapping of RNA-Binding Regions (RBR) in CTCF

To distinguish if the observations above were due to a direct disruption of CTCF-RNA interactions or indirect effects of inhibiting transcription, we generated RNA binding-deficient CTCF mutants in mESCs. Given that homozygous deletion of CTCF is embryonically lethal (Kemp et al., 2014; Moore et al., 2012), we induced the rapid degradation of endogenous CTCF using the auxin-inducible degron system (Nora et al., 2017); degradation was lethal after 2-4 days. To bypass this issue, we transduced cells with lentivirus containing a vector encoding an internal ribosomal entry site (IRES) that allows a wild-type or mutant version of CTCF and the red fluorescent protein mCherry to be simultaneously expressed from a single mRNA transcript. We then sorted the successfully infected cells to obtain a pooled population of steady-state rescues after degradation of the endogenous CTCF protein via the auxin-inducible degron (**Figure S2A**).

To generate defined RNA binding-deficient mutants, we focused on two regions detected by RBR-ID (He et al., 2016): one overlapping part of ZF1 (aa 264-275; KTFQCELCSYTCPR) and another within ZF10 (aa 536-544; QLLDMHFKR), the latter identified within our earlier biochemical mapping of CTCF (Saldaña-Meyer et al., 2014) (**Figure 2A**). Henceforward, the deletion of these 14 and 9 amino acids from ZF1 and ZF10, respectively, will be denoted as ZF1∆ and ZF10∆. While the deletions rescued lethality of endogenous CTCF depletion and had comparable levels of expression (**Figure S2B**), the cells exhibited a significantly slower proliferation rate (**Figure S2C**). These results suggested that an important biological role of CTCF involves ZF1 and ZF10.

**Figure 2.**
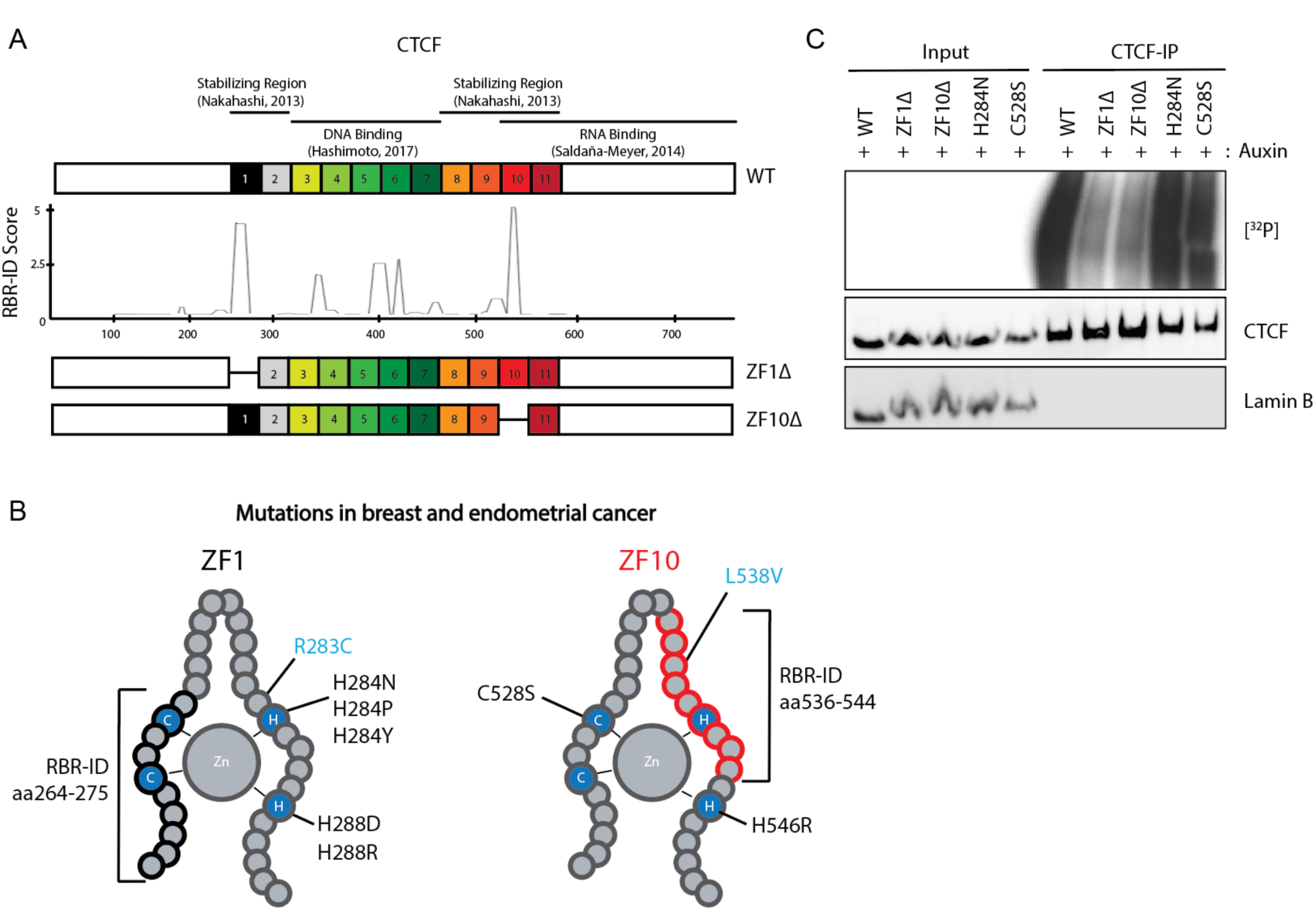
Deletions in ZF1 and ZF10 independently abolish CTCF binding to RNA. **(A)** Schematic representation of known domains of CTCF WT with its 11 zinc fingers being numbered (top); smoothed residue-level RBR-ID score (He et al., 2016) plotted along the primary sequence (middle); and schematics of RBR-deleted constructs used for rescues (bottom). **(B)** Schematic representation of ZF1 and ZF10 of CTCF, mutations found in breast and endometrial cancer that alter ZF binding are shown in black, mutations that do not alter ZF are in blue and RBR-ID deletions are in brackets. **(C)** PAR-CLIP of stably expressed WT and mutant CTCF in mESCs. Autoradiography for ^32^P-labeled RNA (top) and control western blot (middle & bottom).

Notably, we found naturally occurring mutations in endometrial and breast cancer within ZF1 (H284N) and ZF10 (C528S) that target the histidine or cysteine residues that are essential for zinc binding in C_2_H_2_ type ZFs (Kemp et al., 2014)(**Figure 2B**). To test the RNA binding capacity of these mutants, we used photoactivatable ribonucleoside-enhanced cross-linking and immunoprecipitation (PAR-CLIP). The WT rescue showed robust binding to RNA molecules whereas both ZF1∆ and ZF10∆ mutants displayed a drastic reduction in binding as evidenced by a significantly decreased signal in radiolabeled RNA compared to the WT rescue. Surprisingly, point mutations within ZF1 (H284N) or ZF10 (C528S) had no effect on RNA binding (**Figure 2C**). These results suggest that CTCF binding to RNA is not just a consequence of simple RNA affinity to its ZFs, but instead requires a structural conformation that is disrupted independently by the deletion of RBRs in ZF1 and ZF10.

### Deletion of RBRs in CTCF disturb chromatin binding and gene expression

To test if the presence of RNA binding-defective mutants had any effect on gene expression we subjected ZF1∆ and ZF10∆ rescue cell lines to single-cell RNA-seq (scRNA-Seq). Principal component analysis (PCA) showed the similarity between the different rescues, but also underscored the clear distinction between each population of cells (**Figure 3A and S3A**). Further analysis on the differential expression represented as heatmaps showed that genes that are downregulated are similar in both mutant rescues, but there are distinct clusters of genes being upregulated for ZF1∆ and ZF10∆ (**Figure 3B**).

**Figure 3.**
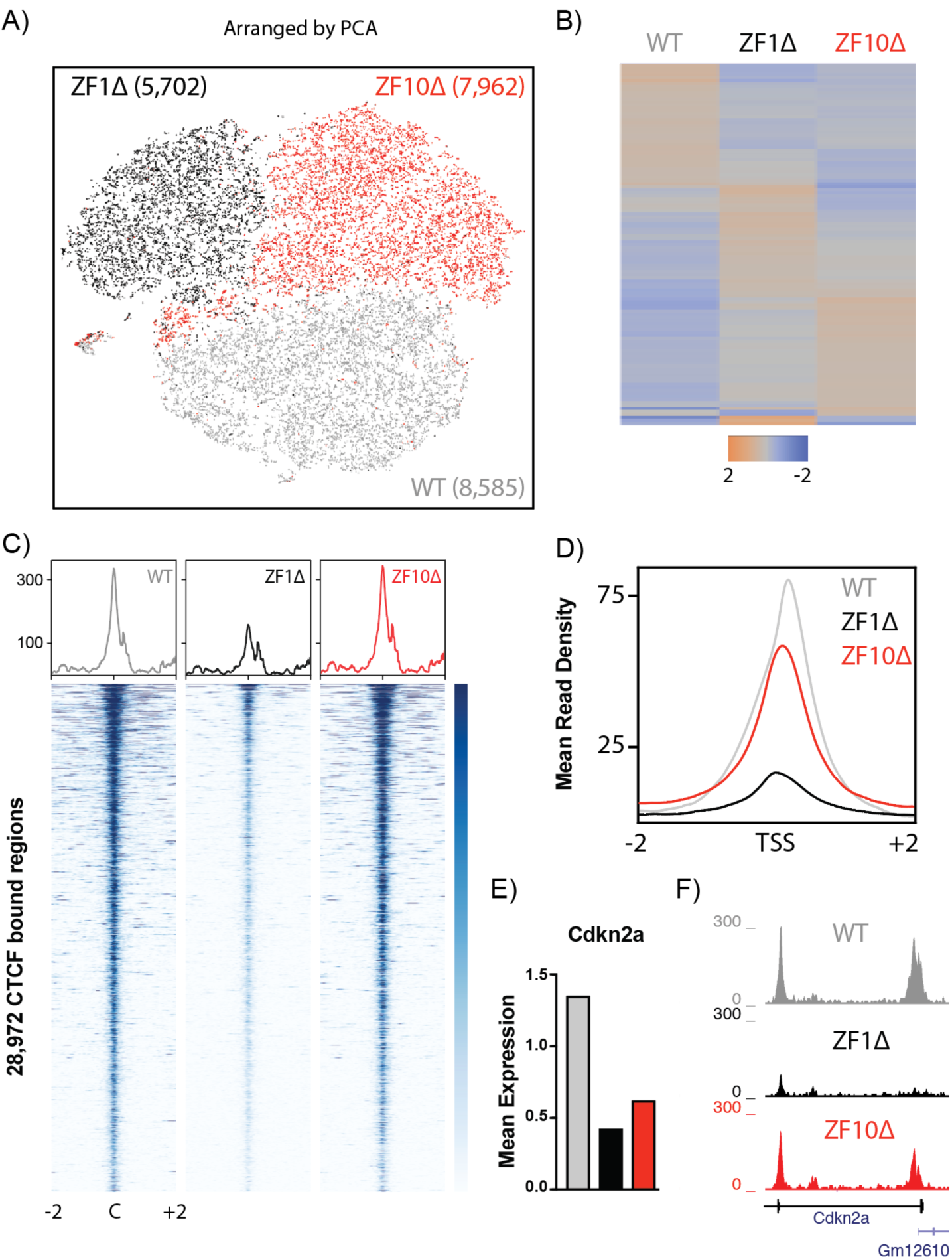
Deletions of RBRs in CTCF cause gene expression defects. **(A)** Principal component analysis (PCA) based representation of single-cell RNA-seq data for rescue cell lines from WT (grey), ZF1∆ (black) and ZF10∆ (red). Each dot represents a single-cell and dots are arranged based on PCA. The final number of cells sequenced per condition is noted in parenthesis. **(B)** Heatmaps depicting differentially expressed genes from scRNA-Seq. **(C)** CTCF ChIP-seq for the same cell lines as in A. Heatmaps were generated by centering and rank-ordering on CTCF binding sites. Corresponding average density profiles are plotted at the top of the heatmaps. **(D)** Average density profile for the ChIP-Seq in (C), but centered in the transcriptional start sites of all genes. **(E)** Mean expression levels for differentially expressed gene *Cdkn2a* and **(F)** corresponding ChIP-Seq tracks.

Furthermore, findings from ChIP-Seq experiments reflected those presented above for TI. The ZF1∆ rescue showed a striking decrease in binding genome-wide while the ZF10∆ rescue seemed to have an overall binding profile that was comparable to the WT rescue (**Figure 3C**). However, when we focused on regulatory regions we were able to observe a subset of genomic regions that have significantly reduced binding of both ZF1∆ as well as ZF10∆ mutants at TSSs and enhancers (**Figure 3D; Figure S3B**). These data indicate that although both ZF1∆ and ZF10∆ have similar deficiencies in RNA binding (**Figure 2B**) they appear to have distinct gene expression and binding profiles.

scRNA-Seq allows us to monitor the variability and consistency of the phenotypes we observe at the single-cell level but is not regularly used to test differential expression due to the lower sequencing depth per cell. Because of these limitations, we also performed regular bulk RNA-seq and compared the two approaches. The differentially expressed genes (DEG) for ZF1∆ and ZF10∆ rescue cell lines showed good overlap (around 50% or more) (**Figure 3E-F**) with each other even at different thresholds of significance and between bulk and scRNA-Seq (**Figure 4A**). Comparing all DEGs from both RNA binding-deficient mutants to all DEGs from cells depleted of CTCF for 24 and 48 hours provided similar overlaps (**Figure 4A**). Remarkably, all these different DEGs displayed a distinctive similarity with nearly all their promoters and/or gene bodies presenting CTCF binding sites, a feature that is significantly lower for genes picked at random (**Figure 4B**). Furthermore, when looking at the motif affinity of those CTCF sites at TSSs and intragenic regions of those DEGs, we observed a significantly lower score versus control sites (**Figure 4C**). Considering the binding profile of CTCF on the TSSs of these DEG we can distinguish at least two groups, one that has CTCF binding enriched at or near TSSs and one without it in the WT situation (**Figure 4D**). Furthermore, upregulated genes after 24 hours of depletion of endogenous CTCF seem to have CTCF binding sites on their TSSs more frequently than those that are downregulated. Such differences were not observed for ZF1∆ and ZF10∆ DEGs (**Figure 4D**). It was previously shown that after 24 hours of CTCF depletion downregulated genes were enriched for CTCF binding sites at TSSs (Nora et al., 2017) yet our results suggest no distinction between down and upregulated genes. These results together suggest that mutating the RBRs in CTCF have effects that partially but not completely overlap with acute depletion of CTCF.

**Figure 4.**
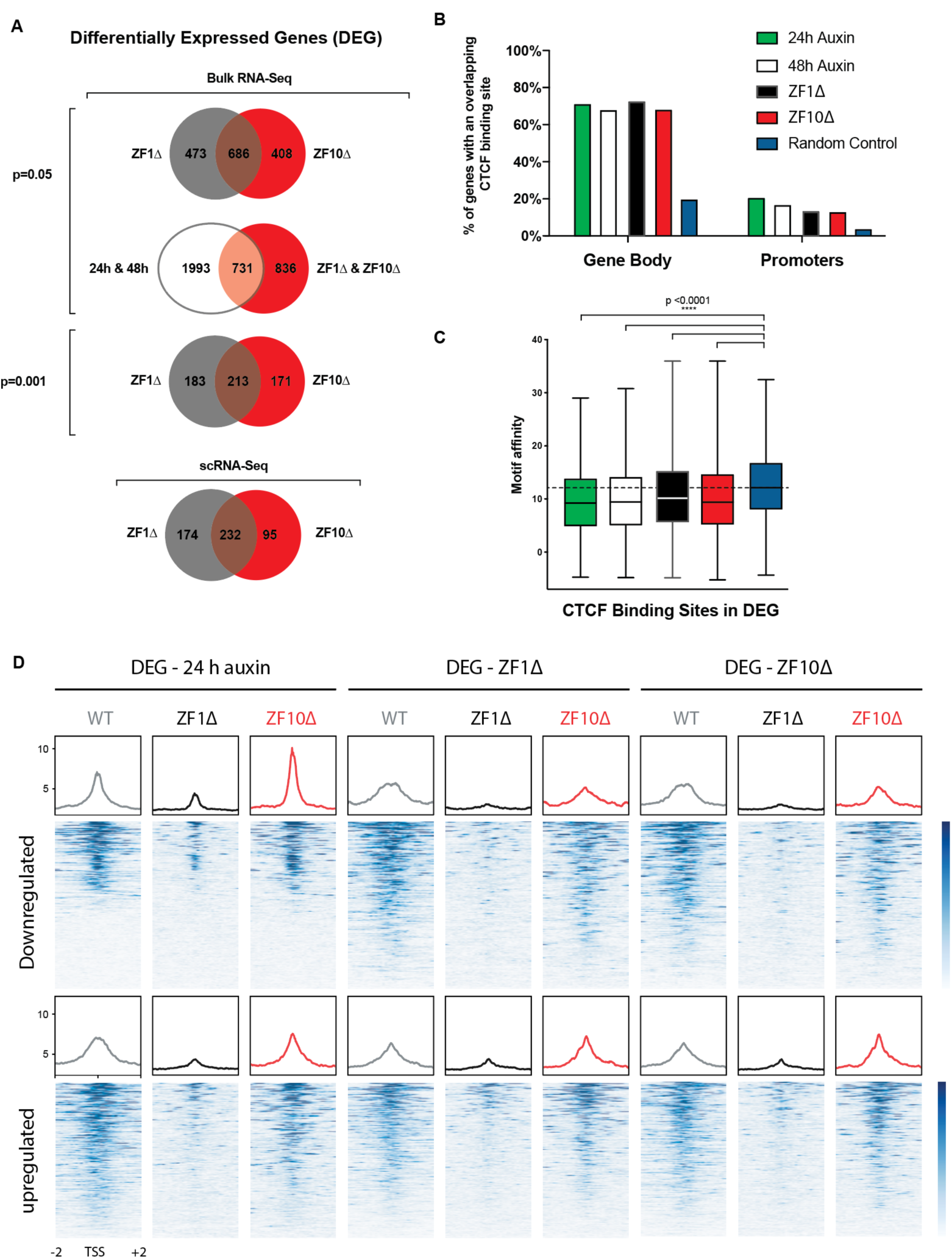
Gene expression defects are partially preserved between conditions. **(A)** Venn diagrams showing the overlap between differentially expressed genes for the different conditions and levels of significance. **(B)** Graph bar illustrating the percentage of genes that have at least one CTCF binding site for CTCF in the promoter region or gene body. **(C)** Box plot showing the motif affinity scores for CTCF binding sites within DEG represented in B compared to a random sample of genes. **(D)** CTCF ChIP-seq for each cell line indicated. Heatmaps were generated by centering and rank-ordering on DEG for each condition shown. Corresponding average density profiles are plotted at the top of the heatmaps.

### Deletion of RBRs in CTCF disturb 3D Chromatin Structure

In order to measure changes in chromosome organization we performed Hi-C experiments on the ZF1∆ and ZF10∆ rescue cell lines. First, we focused on genomic compartmentalization using PCA and hierarchical clustering, which reveals spatial segregation into A “active” and B “inactive” chromatin compartments. Neither RNA binding-deficient mutant rescue showed changes in the plaid pattern, as defined by the eigen-vectors of the Hi-C correlation map nor in compartment domains compared to the WT control (**Figure S4**). This is consistent with previous observations revealing that genomic compartmentalization relies on mechanisms independent of CTCF and cohesin (Nora et al., 2017; Rao et al., 2017).

Next, we focused on the local insulation at TAD boundaries. TADs are highly self-interacting chromatin whose boundaries are enriched with the insulator protein CTCF. The main function of these insulated structures is to restrict the interactions of enhancers with promoters of genes located within the same domain (Rowley and Corces, 2018).

For these analyses we first called TADs for the WT cell line and then used the cut-off value as the benchmark to call TADs on mutant lines. We observed a lower number of TADs called and the median sizes of those TADs increased in both mutants (**Figure S5C-D**). Additionally, we calculated mean boundary scores (MBS) or insulation Scores (IS) (Crane et al., 2015; Lazaris et al., 2017). We observed a significant decrease in insulation of TAD boundaries for both ZF1∆ and ZF10∆ (**Figure 5A-B**). While ZF1∆ has a more general decrease of insulation scores, ZF10∆ shows an intermediate phenotype.

**Figure 5.**
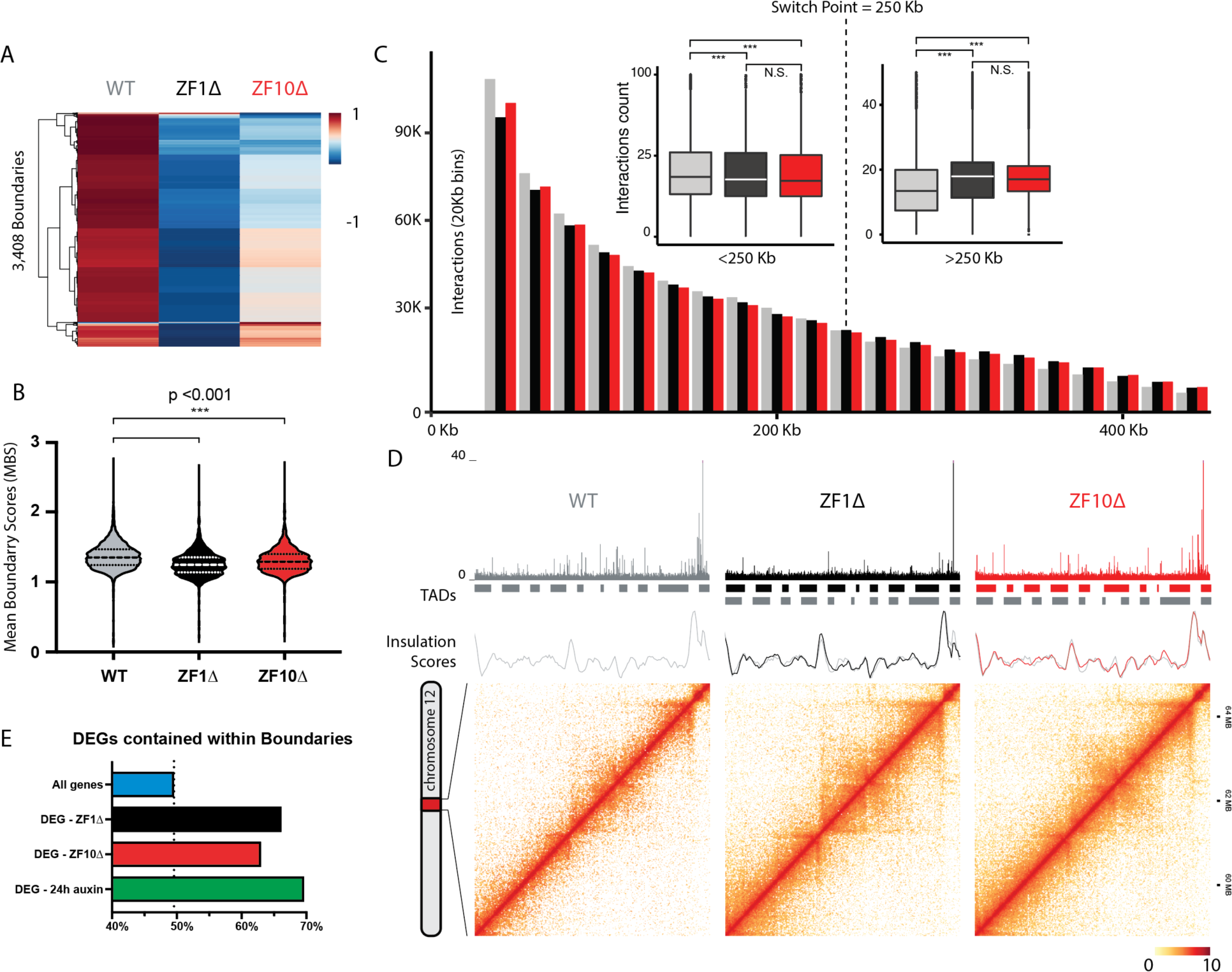
RBR mutants correlates with a global weakening of boundaries. **(A)** Heatmap of Mean Boundary Scores (MBS) across samples. Clustered by row and column. **(B)** Violin plot for MBS, the median value and quartiles are indicated by dotted lines. Statistical significance was assessed using a paired two-sided Wilcoxon rank-sum test (p<0.001). **(C)** Bar-graph showing the decrease of short-range interactions (~40 kb-250 Kb) and increase of long-range interactions (~250-550 Kb) in the mutants. The switch point is represented with dotted lines. **(Left)** Box plot quantifying the differences in interaction distances <250 Kb and **(Right)** >250 Kb across samples. Statistical significance was assessed using a paired two-sided Wilcoxon rank-sum test (p<0.001). **(D)** Hi-C contact maps at 20 kb resolution for a representative region in chromosome 12. From top to bottom, ChIP-Seq tracks for CTCF are shown, then TADs and finally insulation scores graphed along the region. **(E)** Bar graph showing the percentage of DEGs contained within the regions defined as boundaries. DEGs for ZF1∆, ZF10∆ and 24 h of acute depletion of CTCF show a higher localization to boundary regions. All genes contained within the same boundaries serve as a control.

Whereas boundaries have decreased insulation, the overall far *cis*-interactions were significantly increased. Both, ZF1∆ and ZF10∆ decrease in interactions lower than 250 Kb but increase in the range of ~250-550 Kb, shifting their respective median values up from 2,820 Kb for WT to 3,430 Kb for ZF1∆ and 3,900 Kb for ZF10∆ (**Figure 5C**). These differences are further evidenced by Hi-C contact maps (**Figure 5D; Figure S5A-B**).

Integrating ChIP-Seq and Hi-C contact maps, we can appreciate the distinct contributions of each mutant. The overall binding to chromatin is decreased for ZF1∆ where local insulation decreases and new far *cis*-interactions arise and the binding of ZF10∆ is largely conserved but the structural phenotype remains (**Figure 5D; Figure S5A-B**). Finally, we analyzed boundary regions that showed lower insulation and asked if there was any correlation with DEGs. We quantified the percentage of genes contained within regions defined as TAD boundaries. All genes served as a control for the natural distribution of genes within boundaries. DEGs for ZF1∆, ZF10∆ and 24 h after acute depletion of CTCF showed a higher localization to boundary regions compared to all genes (**Figure 5F**). These results suggest that the interactions between CTCF, RNA and chromatin are required for proper local insulation and gene expression.

## Discussion

We previously demonstrated the ability of CTCF to bind large numbers of RNAs (Saldaña-Meyer et al., 2014) and these findings were corroborated by others (Kung et al., 2015). In this study, we were able to dissect a fundamental and general role for RNA-binding to CTCF. Importantly, we describe a clear co-dependency of proper binding to RNA and chromatin that in turn directly disturbs transcription and 3D chromatin structure.

In this study, we provide new insights into the relevance of CTCF-RNA interactions. We demonstrate that local insulation requires not only CTCF binding to chromatin but also to RNA. The reduced insulation observed in RNA binding-deficient mutants causes an overall increase in far *cis*-interactions highlighting the need for RNA molecules to stabilize and demarcate boundaries. Specifically, both ZF1∆ and ZF10∆ show a decrease in interactions lower than 250 Kb but an increase in the range of ~250-550 Kb (**Figure 5C**). This is particularly relevant since it suggests that the insulation of intra-TADs is more sensitive to disruption, clarifying why the size of TADs increases in RNA binding-deficient mutants (**Figure S5B**). These results suggest that new long-range *cis*-interactions that are formed throughout the genome alter cell proliferation (**Figure S2B**). Furthermore, gene expression alterations are enriched at boundary regions (**Figure 3 and 4)**, possibly through new or disrupted promoter-enhancer contacts or aberrant inter-TAD interactions suggesting these are important regulatory regions.

We previously hypothesized that RNA molecules would stabilize CTCF-CTCF loops *in vivo* after describing that CTCF self-association was RNA-dependent *in vitro* (Saldaña-Meyer et al., 2014). In our previous study, we termed the RBR as the fragment of CTCF from ZF10 to the end of the C-terminus and a deletion within that RBR (aa574-614) was observed to be necessary for CTCF self-association and affected RNA binding *in vitro* (Saldaña-Meyer et al., 2014). Unexpectedly, in contrast to in vitro results when we performed PAR-CLIP on a full-length CTCF with the internal deletion we only observed a modest reduction in RNA binding (data not shown) compelling us to pursue further mapping of RBRs that are now presented in this study.

Some examples exist of individual RNAs that have important and specific functions and we expect more will surface in the future, especially for different cell types or during specific stages of development and differentiation. Regardless, we favor the view that most RNA molecules, not only ncRNAs, have a structural and stabilizing role inside the nucleus, as well as the potential to mediate or increase protein-protein interactions without showing any obvious sequence-specificity.

The concept of RNA as a structural component of the nucleus originated in 1989 when the Sheldon Penman group reported that the nuclear matrix fibers collapse and aggregate after treatment with RNase A or actinomycin D in detergent-extracted cells. They proposed that RNA is a structural component of the nuclear matrix, which in turn might organize the higher order structure of chromatin (Nickerson et al., 1989). More recently, Hall *et al.* showed that RNAs transcribed from repetitive LINE1 elements stably associate with interphase chromosomes and are stable under transcriptional inhibition. Furthermore, the loss of these nuclear RNAs from euchromatin disrupts proper chromatin condensation underscoring the putative structural role for transposons, including LINEs and other repetitive sequences that together comprise more than half of the human genome (de Koning et al., 2011; Hall et al., 2014).

CTCF is highly conserved across species (Heger et al., 2012; Rudan et al., 2015), and the presence of its eleven ZFs suggest that it can bind DNA in multiple ways (Filippova et al., 1996; Nakahashi et al., 2013). The 20-base-pair DNA core motif (Holohan et al., 2007; Kim et al., 2007; Schmidt et al., 2010; Xie et al., 2007) was suggested to be engaged by ZFs 4–7 *in vivo* (Nakahashi et al., 2013). This motif is present in most of the known CTCF-binding sites identified by ChIP-seq, and the nonspecific engagement of ZFs other than 4–7 with the flanking DNA sequence was proposed to stabilize CTCF binding (Nakahashi et al., 2013). Recently, the crystal structure of overlapping stretches of its ZFs bound to its core motif was resolved showing that ZF3-7 engage the mayor groove of the core DNA motif. Importantly, it also revealed the lack of a specific function in DNA recognition/binding for ZF1, ZF10 and ZF11 (Hashimoto et al., 2017). Furthermore, mutating the histidine of ZF1 was previously shown to modestly affect the binding of CTCF to chromatin (Nakahashi et al., 2013) but our results show that a comparable point mutation did not affect RNA binding however a deletion within ZF1 had a significant decrease in both RNA and chromatin binding (**Figure 2B-C**). Lastly, point mutations in ZF1 and 10 that do and do not affect the binding of ZFs are found in cancer (**Figure 2B**) (Kemp et al., 2014). In the context of this study, these data together suggest that the main property of ZF1 and ZF10 is binding to RNA rather than DNA.

Thus far, CTCF binding to DNA seems unaffected by other factors as the knockdown of most of its binding partners is ineffectual. One exception is the general transcription factor II-I (TFII-I) that seems to stabilize CTCF binding at promoter regions (Peña-Hernández et al., 2015). In the context of our observations under TI (**Figure 1**), the knockdown of TFII-I most likely affects the transcription of its target genes and hence the decrease in CTCF binding might be an indirect effect of disrupting transcription. Here, we showed that the CTCF binding sites mostly affected are those whose sequence diverged significantly from its core DNA-binding motif (**Figure 1D and 4C**), suggesting that these variations can have important roles in regulatory mechanisms with RNA-binding at its core. Determining how CTCF interacts in complex with DNA and RNA as well as with its protein partners will be an exciting new research avenue.

CTCF was originally described as a transcription factor and there are several examples showing that CTCF binding to gene promoters is necessary for proper transcription of tumor suppressor genes such as BRCA1, RB, TP53 and p16INK4a (Butcher and Rodenhiser, 2007; La Rosa-Velazquez et al., 2007; Soto-Reyes and Recillas-Targa, 2010; Witcher and Emerson, 2009). Yet, since the description of CTCF role as an architectural protein, little attention has been afforded to its role as a transcription factor. Most arguments against CTCF being important for gene expression rest on the relatively small number (~200-400) of genes that are affected upon its knockdown or even its acute depletion using an auxin-inducible degron (Nora et al., 2017; Zuin et al., 2014). In this context, our ZF1∆ and ZF10∆ rescue cell lines also had modest numbers of DEGs, although this varied depending on the threshold applied ~1000 genes (adjusted p<0.05) to ~400 (adjusted p<0.001).

Additionally, when analyzing the relative occupancy of CTCF measured by ChIP-Seq, promoter regions have significantly less occupancy compared to the overall binding sites (**Figure 3A and D**) (Weintraub et al., 2017). CTCF binding sites with low occupancy that diverge from the core DNA motif were described to be associated with regulated binding during mESC differentiation (Plasschaert et al., 2014). In our study, we noticed that CTCF binding sites within DEGs have the same characteristics and CTCF binding to them is destabilized in ZF1∆ and ZF10∆ rescue cell lines compared to WT (**Figure 4C**).

It is our view that CTCF has a significant role in regulating gene expression based on the following observations: 1) hemizygous mice for CTCF succumb to cancer in 80% of animals tested. Highlighting that chronically lower levels of CTCF have clear dramatic effects on the biology of the cell and seem to be a hallmark of carcinogenesis (Kemp et al., 2014), 2) differentially expressed genes have significantly more CTCF binding sites in their promoters and gene bodies (**Figure 4B**), 3) CTCF orientation at promoters is in the same direction as transcription and these form loops with internal CTCF binding sites close to exons. These loops are prevalent and significant for alternative splicing (Ruiz-Velasco et al., 2017), 4) differences between cell-type specificity of CTCF binding is fitting with the transcription trapping hypothesis: RNA contributes to the maintenance and recognition of its binding site for certain transcription factors such that “transcription of regulatory elements produces a positive-feedback loop that contributes to the stability of gene expression programs” (Sigova et al., 2015) and 5) highly expressed genes can sometimes displace CTCF and push cohesin to intergenic CTCF sites where they can form chromatin loops (Busslinger et al., 2017; Heinz et al., 2018). All these features together underscore the relevance of CTCF as a transcription factor and the interplay between transcription and chromatin organization.

Many questions remain to be explored if transcription is considered to be a main factor contributing to the regulation of TADs in a temporal and cell-type specific manner. Perhaps other RNA binding proteins can account for specific structural roles. Many TADs are either gene poor or the genes they contained are largely silenced, in these cases there can conceivably be no contribution of RNA as a structural component. However, it is possible that abundant long-lived transcripts such as those from repetitive regions (Hall et al., 2014) could have a general function in chromatin organization of these repressed regions. Based on the large number of RNA interactors that are pulled-down with CTCF (Kung et al., 2015; Saldaña-Meyer et al., 2014) we envision these interactions might be highly redundant and technical breakthroughs will be needed to improve our understanding of these complex regulatory mechanisms.

## ACKNOWLEDGMENTS

We thank Drs. L. Vales for comments on the manuscript; Dr. Roberto Bonasio for sharing pre-publication results for RBR-ID; Dr. Esteban Mazzoni for continuing discussions; Dr. Robert Tjian and collaborators for early access to their manuscript; New York University Langone Medical Center (NYULMC) Genome Technology Center; NYULMC Cytometry and Cell Sorting Core for help with FACS; Programa de Apoyo a Proyectos de Investigación e Innovación Tecnológica (PAPIIT, IA201817 and IN207319) for financial support and the Laboratorio Nacional de Ciencias de la Sostenibilidad (LANCIS, UNAM) and Ing. Rodrigo García Herrera for access and technical support with the computer cluster.

The NYULMC Genome Technology Center and the NYUMC Cytometry and Cell Sorting Core are partially supported by the Cancer Center Support Grant, P30CA016087, at the Laura and Isaac Perlmutter Cancer Center. This work utilized computing resources at the High-Performance Computing Facility of the Center for Health Informatics and Bioinformatics at the NYULMC.

This work was supported by grants from the NIH (1R01NS100897 to D.R.), R35GM122515 and 1R01-CA222131 (to J.S.) and the Howard Hughes Medical Institute (to D.R.).

## AUTHOR CONTRIBUTIONS

R.S.M and D.R. conceived the project, designed the experiments, and wrote the paper with input from all authors; R.S.M. performed all experiments and bioinformatic data analysis; E.N. generated the CTCF degron cell lines in the lab of B.B.; J.R.H. and M.N. performed bioinformatic analysis for Hi-C data in the lab of J.S.; K.J.L. analyzed Hi-C data in the lab of M.F.M.

## DECLARATION OF INTERESTS

D.R. is a co-founder of Constellation Pharmaceuticals and Fulcrum Therapeutics. All other authors declare that they have no competing interests.

## STAR METHODS

## KEY RESOURCES TABLE

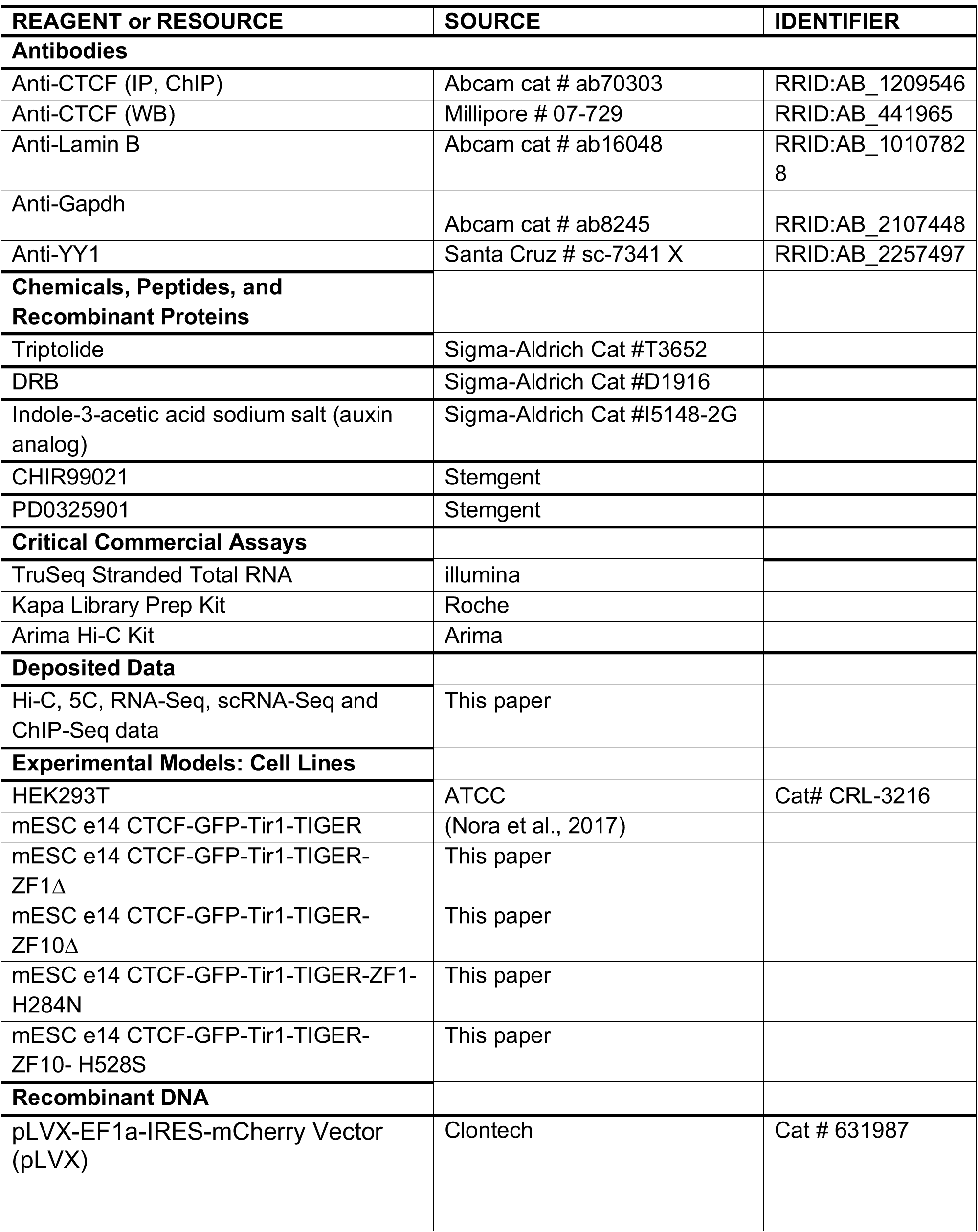
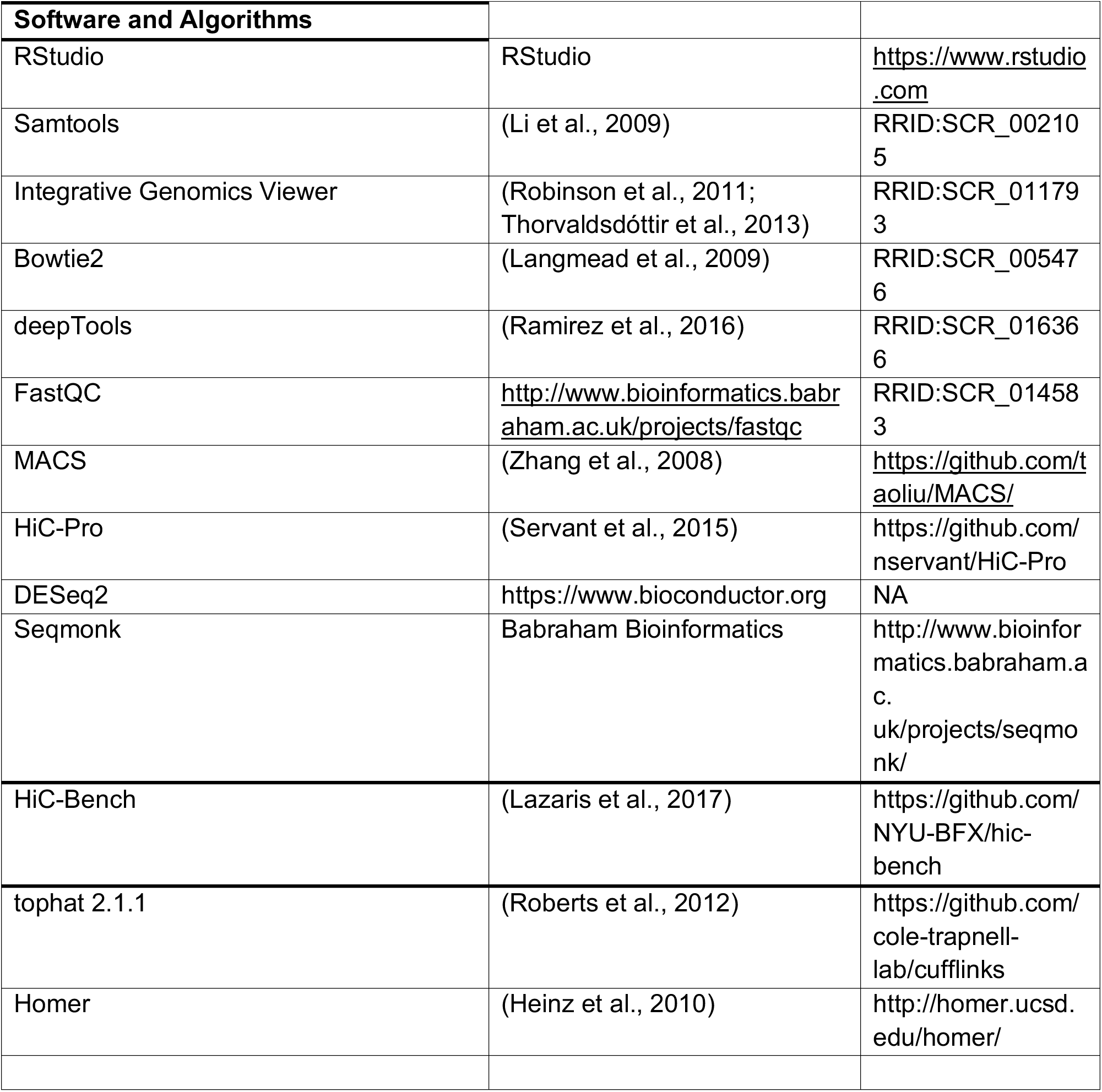

## CONTACT FOR REAGENT AND RESOURCE SHARING

Further information and requests for resources and reagents should be directed to and will be fulfilled by the Lead Contact, Danny Reinberg

## EXPERIMENTAL MODEL AND SUBJECT DETAILS

### Mouse ESC culture

E14Tga2 (ATCC, CRL-1821) mESCs were grown in standard medium supplemented with LIF, 1 mM MEK1/2 inhibitor (PD0325901, Stemgent) and 3 mM GSK3 inhibitor (CHIR99021, Stemgent); rescue cell lines were also grown with 500mM of indole-3-acetic acid (IAA, chemical analog of auxin) in the medium as in (Nora et al., 2017).

## METHOD DETAILS

### Transcriptional inhibition

mESCs were incubated with a combination of transcriptional inhibitors (Triptolide 1 µM and DRB 100µM) for 1 or 4 hours. After treatment cells were immediately harvested for immunoblot, ChIP and 5C experiments.

### RNase A treatment

The treatment was performed as in (Beltran et al., 2016). Briefly, mESCs nuclei were permeabilized with 0.05% Tween-20 in PBS for 10 min at 4C, washed once and resuspended in PBS and finally incubated with RNase A (1 mg/ml) or a mock reaction for 30 minutes. After treatment cells were immediately harvested for ChIP and 5C experiments.

### Rescue cell line generation

HEK293T to cells were grown to ~90% confluency, split 1:4 and grown for one day. Cell were then transfected with pLVX-EF1a-IRES-mCherry encoding CTCF WT, ZF1∆, ZF10∆, H284N or C528S with their respective packaging vectors. After 4 hours of transfection the medium was changed to complete DMEM and finally to ESC culture medium after 32-40hr of transfection. Then, the viral supernatant was harvested, filtered through 0.45um syringe filter and added polybrene to 8µg/µl. Added the mix to cells and spin infect (500g X 60min). Change medium the next day. Sorted for mCherry positive cells after ~2 days of infection for each condition.

### PAR-CLIP

PAR-CLIP was performed as in (Saldaña-Meyer et al., 2014) with some modifications. Briefly, cells were grown under standard conditions and pulsed with 400 mM 4-SU (Sigma) for 2 h. After washing the plates with PBS, cells were cross-linked with 400 mJ/cm2 UVA (312 nm) using a Stratalinker UV cross-linker (Stratagene). Whole nuclear lysates (WNLs) were obtained by fractionated and nuclei were then incubated for 10 min at 37°C in an appropriate volume of CLIP buffer (20 mM HEPES at pH 7.4, 5 mM EDTA, 150 mM NaCl, 2% lauryldimeth-ylbetaine) supplemented with protease inhibitors, 20 U/mL Turbo DNase (Life technologies), and 200 U/mL murine RNase inhibitor (New England Biolabs). After clearing the lysate by centrifugation, immunoprecipitations were carried out using 200 µg of WNLs, CTCF antibody, in the same CLIP buffer for 4h at 4°C and then added protein G-coupled Dynabeads (Life Technologies) for an additional hour. Contaminating DNA was removed by treating the beads with Turbo DNase (2 U in 20 mL). Cross-linked RNA was labeled by successive incubation with 5 U of Antarctic phosphatase (New England Biolabs) and 5 U of T4 PNK (New England Biolabs) in the presence of 10 mCi [g-32P] ATP (PerkinElmer). Labeled material was resolved on 8% Bis-Tris gels, transferred to nitrocellulose membranes, and visualized by autoradiography. The same membrane was then blocked with TBS-T and 5% milk and blotted for CTCF and Lamin-B.

### ChIP-seq

ChIP-seq experiments were performed as described previously (Gao et al., 2012). Briefly, cells were fixed with 1% Formaldehyde. Nuclei were isolated using buffers in the following order: LB1 (50 mM HEPES, pH 7.5 at 4C, 140 mM NaCl, 1 mM EDTA, 10% Gly- cerol, 0.5% NP40, 0.25% Triton X; 10 min at 4C), LB2 (10 mM Tris, pH 8 at 4C, 200 mM NaCl, 1 mM EDTA, 0.5 mM EGTA; 10 min at RT), and LB3 (10 mM Tris, pH 7.5 at 4C, 1 mM EDTA, 0.5 mM EGTA, and 0.5% N-Lauroylsarcosine sodium salt). Chromatin was fragmented to an average size of 250 bp using a Diagenode Bioruptor. Chromatin immunoprecipitation was performed with CTCF or K27ac antibodies. Chromatin from Drosophila (in a 1:50 ratio to the ESC-derived chromatin) as well as Drosophila specific H2Av antibody was used as spike-in control in rescue cell lines. ChIP-seq libraries were prepared using the Kapa Library Prep Kit.

### Bulk RNA-seq

Total RNA from ESCs was isolated with TRIzol (Life Technologies). Stranded libraries were prepared using TruSeq Stranded Total RNA kits.

### Single-cell RNA-seq library preparation

Single-cell RNA-seq libraries were prepared using the Chromium Controller (10X Genomics). Briefly, single cells in 0.04% BSA in PBS were separated into droplets and then reverse transcription and library construction was performed according to the 10X Chromium Single Cell 30 Reagent Kit User Guide and sequenced on an Illumina Novaseq 6000.

### Hi-C library preparation

Hi-C libraries were constructed using the Arima kit following the manufacturer’s instructions.

### Definition of regulatory regions

### Promoters

For Figure S1D promoters were defined as ± 5 kilobases from the transcription start site.

### Enhancers

For Figure S1C and S3B. Typical-enhancer coordinates were downloaded from (Whyte et al., 2013)

### 5C Library Preparation

5C was performed as in (Narendra et al., 2016). Briefly, 5C primers were annealed at 48°C for 16hrs atop the 3C libraries from each sample. 1fmol of each primer was used in the annealing reaction with 1μg of 3C template and 1μg of salmon sperm DNA. 16 separate annealing reactions were performed per sample, along with control reactions with individual components removed. Forward and reverse primers that annealed to adjacent regions of the 3C template were ligated with 10U of Taq ligase for 60’ at 48°C. Successfully ligated forward-reverse primer pairs were then amplified in 6 separate PCR reactions per annealing reaction, using primers specific to the T7 and T3 overhangs. PCR reactions from the equivalent initial sample were then pooled, purified and run on a gel to ensure the control reactions did not show an amplification product. Libraries were then generated from the purified PCR product to allow for deep sequencing.

## QUANTIFICATION AND STATISTICAL ANALYSIS

### Bulk RNA-seq Analysis

Raw sequencing files were aligned against the mouse reference genome (GRCm38/mm10) using tophat 2.1.1 and differentially expressed genes were called using DESeq2 with adjusted p-values 0.05 and 0.001.

### Single-cell RNA-seq Analysis

Sequencing data was demultiplexed using the 10X Genomics Cell Ranger software (version 2.0.0) and aligned to the mm10 transcriptome. Unique molecular identifiers were collapsed into a gene-barcode matrix representing the counts of molecules per cell as determined and filtered by Cell Ranger using default parameters. Normalized expression values were generated using Cell Ranger using the default parameters.

### ChIP-seq Analysis

Raw sequencing files were aligned against the mouse reference genome (GRCm38/mm10) using Bowtie2 (version 2.3.4.1) (parameters: -N 1 -k 1 -q -x). Ambiguous reads were filtered to use uniquely mapped reads in the downstream analysis. PCR duplicates were removed using Picard-tools (version 1.88). MACS version 2.0.10 was used to call narrow peaks (parameters: -g 1.87e9 –qvalue 0.01). To create heatmaps we used deepTools (version 2.4.1) (Ramirez et al., 2016). We first ran bamCoverage (–binSize 50 –extendReads 200 -of bigwig) and normalized read numbers to RPKM or to the spike-in drosophila DNA (–scaleFactor sf), obtaining read coverage per 50-bp bins across the whole genome (bigWig files). We then used the bigWig files to compute read numbers centered on CTCF peaks called by MACS, on TSSs or enhancers (computeMatrix reference-point –sortRegions descend –sortUsing mean –averageTypeBins mean). Finally, heatmaps were created with plotHeatmap (–colorMap=’Blues’ –sortRegions=no).

### Hi-C data processing and quality control

#### Processing

HiC-Bench (Lazaris et al., 2017) was used to align and filter the Hi-C data, identify TADs, and generate Hi-C heatmaps. To generate Hi-C filtered contact matrices, the Hi-C reads were aligned against the mouse reference genome (GRCm38/mm10) by bowtie2 (Langmead and Salzberg, 2012) (version 2.3.1). Mapped read pairs were filtered by the Genomic Tools (Tsirigos et al., 2012) tools-hic filter command integrated in HiC-bench for known artifacts of the Hi-C protocol. The filtered reads include multi-mapped reads (‘multihit’), read-pairs with only one mappable read (‘single sided’), duplicated read-pairs (‘ds.duplicate’), low mapping quality reads (MAPQ < 20), read-pairs resulting from self-ligated fragments, and short-range interactions resulting from read-pairs aligning within 25kb (‘ds.filtered’). For the downstream analyses, all the accepted intra-chromosomal read-pairs (‘ds.accepted intra’) were used.

The Hi-C filtered contact matrices were corrected using the ICE “correction” algorithm (Imakaev et al., 2012) built into HiC-bench. TADs and boundaries were called using Crane insulation scores (Crane et al., 2015) (‘ratio’ index) at 40 kb bin resolution with an insulating window of 500kb. HiC heatmaps for regions of interest were generated using the ICE corrected contact matrices through the ‘hic-plotter-diff’ pipeline step integrated in HiC-bench.

Additionally, fastq files were also analyzed using the HiCPro 2.11.1 pipeline (Servant et al., 2015). Mapping was done against the mm10 genome with Bowtie2 (v 2.3.1). Using the same number of valid pairs for all samples we identified TADs using the Insulation Score (IS) metric implemented in TADtool (Kruse et al., 2016) using a resolution of 40kb bins and a window given by default at that resolution of 102,353 bp. The cutoff value for boundary identification through IS, was adjusted using the WT sample (cutoff=50) and applied to all samples (WT, ZF1 and ZF10) for comparison.

### Quality Control and TAD/Boundary Stats

Quality assessment analysis shows that the total numbers of reads in the 2 biological replicates once merged for each condition ranged from ~430 million reads to ~550 million. The percentage of reads aligned was always over 98% in all samples. The proportion of accepted reads (‘ds-accepted-intra’ and ‘ds-accepted-inter’) ranged from ~20% to ~33%, which in all cases was sufficient to call TADs.

## DOWNSTREAM ANALYSIS

### Screening of Potentially Altered Boundaries

The Hi-C downstream analysis involved a global (genome-wide) screening of TAD boundary insulation changes in ZF1 and ZF10 mutants based on boundary insulation scores and CTCF binding enrichment. Individual cases of disrupted boundaries were confirmed by visual inspection of HiC heatmaps.

### Mean Boundary Scores

To assess and compare boundary strength alteration in the ZF mutants, we included the calculation of the mean boundary score (MBS) for every boundary identified in the WT condition (reference boundaries).

The MBS is the arithmetic mean of the insulation scores inside the reference boundary coordinates being assessed. Using the Crane method there is one insulation score per bin (40kb) and as a result there are generally multiple insulation scores per boundary. The MBS was used to calculate ZF1 and ZF2 logFC values with respect to the WT. A differential analysis on the MBS logFC obtained was also performed. An unpaired t-test (two-sided) was used by pooling all the insulation scores inside the boundary and adjusting with FDR correction.

The MBS on R was used to generate heatmaps (‘pheatmap’ function with ‘ward.D2’ clustering method and ‘manhattan’ clustering distance parameters). Boxplots were generated to demonstrate the association of ZF1 and ZF10 mutations with decreased levels of mean boundary scores (MBS). Statistical significance was assessed using a paired two-sided Wilcoxon rank-sum test.

### CTCF Occupancy in Boundaries. Integration with MBS Data

The CTCF peaks were mapped to the boundaries to merge the MBS information obtained with the CTCF binding data. We assigned a peak to a boundary if the peak overlapped with the boundary region (> 0 bp). An extension of the boundary region by 2 bins (80 kb) on either side of the boundary was considered to account for false boundary calls. The CTCF signal of all the peaks assigned to a boundary were aggregated and then ZF1 and ZF10 logFC values were calculated. Significant changes in global CTCF occupancy within the boundaries were calculated using a two-sided unpaired t-test by pooling all the CTCF peaks assigned in the boundary.

A ranked-table of boundary coordinates with the MBS and CTCF metrics was created. To generate Hi-C heatmaps, the best boundary cases were selected by filtering boundaries with MBS FDR > 0.01. We used cutoff values of MBS log2FC < −0.15 and CTCF log2FC < −0.15 to extract boundaries with the strongest MBS / CTCF positive correlation and negative direction. Finally, the remaining boundaries were ranked by MBS logFC and CTCF logFC to obtain a set of affected boundaries based on strong MBS and CTCF occupancy evidence.

### Compartments

Compartment analysis was carried out using the Homer pipeline (Heinz et al., 2010) (v4.6). Homer performs a principal component analysis of the normalized interaction matrices and uses the PCA1 component to predict regions of active (A compartments) and inactive chromatin (B compartments), as well as to identify significantly altered compartments. Homer works under the assumption that gene-rich regions with active chromatin marks have similar PC1 values, while gene deserts show differing PC1 values.

HiC filtered matrices were given as input to Homer together with H2K27ac peaks for compartment prediction. H2K27ac was used by Homer as prior information of active regions.

### DNA-DNA interactions range analysis

An analysis of distance-range of the dna-dna interactions across samples was performed using the ‘interactions’ data produced by HiC-Bench. The interactions are defined in a ‘locus1 vs locus2’ fashion. Every chromosome is divided in bins of 20 kb (locus) where all locus-locus combinations inside a cutoff distance of 10 Mb are assessed. All the counts from the hic-filtered contact matrix are mapped and assigned to a unique locus1-locus2 ID and the total of counts assigned to a locus-1-locus2 ID are aggregated.

We normalized the interaction counts value of every locus1-locus2 ID dividing by the total number of counts obtained in each sample. A table of normalized interaction counts per absolute distance was generated (20kb bins in the range of 40 kb −10 Mb). All the values were multiplied by a scale-factor (Sf) to obtain positive count values higher or equal to 1. The scale-factor was calculated as 1 divided by the minimum normalized value found across all samples (Sf =1/min(all normalized values)). A graph-chart and box-plots was generated on R to show the decrease of short-range interactions (~40kb-250kb) and increase of long-range interactions (250kb-10Mb) in the ZF mutants. To assess statistical significance the counts of each specific distance-bin locus was tested across all the samples by using a Wilcoxon paired one-sided test. This test was performed separately on the loci with a distance smaller than 250kb, and on the loci in the range of 250-10Mb

### Data availability

Data can be found at GEO with accession #XXX

## References

Beltran, M., Yates, C.M., Skalska, L., Dawson, M., Reis, F.P., Viiri, K., Fisher, C.L., Sibley, C.R., Foster, B.M., Bartke, T., et al. (2016). The interaction of PRC2 with RNA or chromatin is mutually antagonistic. Genome Research gr.197632.115.

Crane, E., Bian, Q., McCord, R.P., Lajoie, B.R., Wheeler, B.S., Ralston, E.J., Uzawa, S., Dekker, J., and Meyer, B.J. (2015). Condensin-driven remodelling of X chromosome topology during dosage compensation. Nature 523, 240–244.

Heinz, S., Benner, C., Spann, N., Bertolino, E., Lin, Y.C., Laslo, P., Cheng, J.X., Murre, C., Singh, H., and Glass, C.K. (2010). Simple combinations of lineage-determining transcription factors prime cis-regulatory elements required for macrophage and B cell identities. Molecular Cell 38, 576–589.

Imakaev, M., Fudenberg, G., McCord, R.P., Naumova, N., Goloborodko, A., Lajoie, B.R., Dekker, J., and Mirny, L.A. (2012). Iterative correction of Hi-C data reveals hallmarks of chromosome organization. Nature Methods 9, 999–1003.

Langmead, B., and Salzberg, S.L. (2012). Fast gapped-read alignment with Bowtie 2. Nature Methods 9, 357–359.

Lazaris, C., Kelly, S., Ntziachristos, P., Aifantis, I., and Tsirigos, A. (2017). HiC-bench: comprehensive and reproducible Hi-C data analysis designed for parameter exploration and benchmarking. BMC Genomics 18, 22.

Li, H., Handsaker, B., Wysoker, A., Fennell, T., Ruan, J., Homer, N., Marth, G., Abecasis, G., Durbin, R., 1000 Genome Project Data Processing Subgroup (2009). The Sequence Alignment/Map format and SAMtools. Bioinformatics 25, 2078–2079.

Narendra, V., Bulajić, M., Dekker, J., Mazzoni, E.O., and Reinberg, D. (2016). CTCF-mediated topological boundaries during development foster appropriate gene regulation. Genes & Development 30, 2657–2662.

Roberts, A., Goff, L., Pertea, G., Kim, D., Kelley, D.R., Pimentel, H., Salzberg, S.L., Rinn, J.L., Pachter, L., and Trapnell, C. (2012). Differential gene and transcript expression analysis of RNA-seq experiments with TopHat and Cufflinks. Nature Protocols 7, 562–578.

Tsirigos, A., Haiminen, N., Bilal, E., and Utro, F. (2012). GenomicTools: a computational platform for developing high-throughput analytics in genomics. Bioinformatics 28, 282–283.

Bensaude, O. (2014). Inhibiting eukaryotic transcription. Which compound to choose? How to evaluate its activity? Transcription 2, 103–108.

Bonev, B., and Cavalli, G. (2016). Organization and function of the 3D genome. Nature 17, 661–678.

Busslinger, G.A., Stocsits, R.R., van der Lelij, P., Axelsson, E., Tedeschi, A., Galjart, N., and Peters, J.-M. (2017). Cohesin is positioned in mammalian genomes by transcription, CTCF and Wapl. Nature 544, 1–24.

Butcher, D.T., and Rodenhiser, D.I. (2007). Epigenetic inactivation of BRCA1 is associated with aberrant expression of CTCF and DNA methyltransferase (DNMT3B) in some sporadic breast tumours. European Journal of Cancer 43, 210–219.

de Koning, A.P.J., Gu, W., Castoe, T.A., Batzer, M.A., and Pollock, D.D. (2011). Repetitive elements may comprise over two-thirds of the human genome. PLoS Genetics 7, e1002384.

Filippova, G.N., Fagerlie, S., Klenova, E.M., Myers, C., Dehner, Y., Goodwin, G., Neiman, P.E., Collins, S.J., and Lobanenkov, V.V. (1996). An exceptionally conserved transcriptional repressor, CTCF, employs different combinations of zinc fingers to bind diverged promoter sequences of avian and mammalian c-myc oncogenes. Molecular and Cellular Biology 16, 2802–2813.

Fudenberg, G., Abdennur, N., Imakaev, M., Goloborodko, A., and Mirny, L.A. (2017). Emerging Evidence of Chromosome Folding by Loop Extrusion. Cold Spring Harb. Symp. Quant. Biol. 82, 45–55.

Hall, L.L., Carone, D.M., Gomez, A.V., Kolpa, H.J., Byron, M., Mehta, N., Fackelmayer, F.O., and Lawrence, J.B. (2014). Stable C0T-1 Repeat RNA Is Abundant and Is Associated with Euchromatic Interphase Chromosomes. Cell 156, 907–919.

Hashimoto, H., Wang, D., Horton, J.R., Zhang, X., Corces, V.G., and Cheng, X. (2017). Structural Basis for the Versatile and Methylation-Dependent Binding of CTCF to DNA. Molecular Cell 66, 711–720.e713.

He, C., Sidoli, S., Warneford-Thomson, R., Tatomer, D.C., Wilusz, J.E., Garcia, B.A., and Bonasio, R. (2016). High-Resolution Mapping of RNA-Binding Regions in the Nuclear Proteome of Embryonic Stem Cells. Molecular Cell 64, 416–430.

Heger, P., Marin, B., Bartkuhn, M., Schierenberg, E., and Wiehe, T. (2012). The chromatin insulator CTCF and the emergence of metazoan diversity. Proc. Natl. Acad. Sci. U.S.a. 109, 17507–17512.

Heinz, S., Texari, L., Hayes, M.G.B., Urbanowski, M., Chang, M.W., Givarkes, N., Rialdi, A., White, K.M., Albrecht, R.A., Pache, L., et al. (2018). Transcription Elongation Can Affect Genome 3D Structure. Cell 174, 1522–1536.e1522.

Holohan, E.E., Kwong, C., Adryan, B., Bartkuhn, M., Herold, M., Renkawitz, R., Russell, S., and White, R. (2007). CTCF Genomic Binding Sites in Drosophila and the Organisation of the Bithorax Complex. PLoS Genet 3, e112–e112.

Kaneko, S., Bonasio, R., Saldaña-Meyer, R., Yoshida, T., Son, J., Nishino, K., Umezawa, A., and Reinberg, D. (2014a). Interactions between JARID2 and Noncoding RNAs Regulate PRC2 Recruitment to Chromatin. Molecular Cell 53, 290–300.

Kaneko, S., Son, J., Bonasio, R., Shen, S.S., and Reinberg, D. (2014b). Nascent RNA interaction keeps PRC2 activity poised and in check. Genes & Development 28, 1983–1988.

Kemp, C.J., Moore, J.M., Moser, R., Bernard, B., Teater, M., Smith, L.E., Rabaia, N.A., Gurley, K.E., Guinney, J., Busch, S.E., et al. (2014). CTCF haploinsufficiency destabilizes DNA methylation and predisposes to cancer. CellReports 7, 1020–1029.

Kim, T.H., Abdullaev, Z.K., Smith, A.D., Ching, K.A., Loukinov, D.I., Green, R.D., Zhang, M.Q., Lobanenkov, V.V., and Ren, B. (2007). Analysis of the Vertebrate Insulator Protein CTCF-Binding Sites in the Human Genome. Cell 128, 1231–1245.

Kruse, K., Hug, C.B., Hernández-Rodríguez, B., and Vaquerizas, J.M. (2016). TADtool: visual parameter identification for TAD-calling algorithms. Bioinformatics 32, 3190–3192.

Kung, J.T., Kesner, B., An, J.Y., Ahn, J.Y., Cifuentes-Rojas, C., Colognori, D., Jeon, Y., Szanto, A., del Rosario, B.C., Pinter, S.F., et al. (2015). Locus-Specific Targeting to the X Chromosome Revealed by the RNA Interactome of CTCF. Molecular Cell 1–15.

La Rosa-Velazquez, De, I.A., Rincon-Arano, H., Benitez-Bribiesca, L., and Recillas-Targa, F. (2007). Epigenetic Regulation of the Human Retinoblastoma Tumor Suppressor Gene Promoter by CTCF. Cancer Research 67, 2577–2585.

Lai, F., Orom, U.A., Cesaroni, M., Beringer, M., Taatjes, D.J., Blobel, G.A., and Shiekhattar, R. (2013). Activating RNAs associate with Mediator to enhance chromatin architecture and transcription. Nature 494, 497–501.

Li, W., Notani, D., Ma, Q., Tanasa, B., Nunez, E., Chen, A.Y., Merkurjev, D., Zhang, J., Ohgi, K., Song, X., et al. (2013). Functional roles of enhancer RNAs for oestrogen-dependent transcriptional activation. Nature 498, 516–520.

Merkenschlager, M., and Nora, E.P. (2016). CTCF and Cohesin in Genome Folding and Transcriptional Gene Regulation. Annu Rev Genomics Hum Genet 17, 17–43.

Moore, J.M., Rabaia, N.A., Smith, L.E., Fagerlie, S., Gurley, K., Loukinov, D., Disteche, C.M., Collins, S.J., Kemp, C.J., Lobanenkov, V.V., et al. (2012). Loss of Maternal CTCF Is Associated with Peri-Implantation Lethality of Ctcf Null Embryos. PLoS ONE 7, e34915–10.

Nakahashi, H., Kwon, K.-R.K., Resch, W., Vian, L., Dose, M., Stavreva, D., Hakim, O., Pruett, N., Nelson, S., Yamane, A., et al. (2013). A Genome-wide Map of CTCF Multivalency Redefines the CTCF Code. CellReports 3, 1678–1689.

Nickerson, J.A., Krochmalnic, G., Wan, K.M., and Penman, S. (1989). Chromatin architecture and nuclear RNA. Proceedings of the National Academy of Sciences 86, 177–181.

Nora, E.P., Goloborodko, A., Valton, A.-L., Gibcus, J.H., Uebersohn, A., Abdennur, N., Dekker, J., Mirny, L.A., and Bruneau, B.G. (2017). Targeted Degradation of CTCF Decouples Local Insulation of Chromosome Domains from Genomic Compartmentalization. Cell 169, 930–944.e22.

Pekowska, A., Klaus, B., Xiang, W., Severino, J., Daigle, N., Klein, F.A., Oleś, M., Casellas, R., Ellenberg, J., Steinmetz, L.M., et al. (2018). Gain of CTCF-Anchored Chromatin Loops Marks the Exit from Naive Pluripotency. Cell Syst 7, 482–495.e10.

Peña-Hernández, R., Marques, M., Hilmi, K., Zhao, T., Saad, A., Alaoui-Jamali, M.A., del Rincon, S.V., Ashworth, T., Roy, A.L., Emerson, B.M., et al. (2015). Genome-wide targeting of the epigenetic regulatory protein CTCF to gene promoters by the transcription factor TFII-I. Proceedings of the National Academy of Sciences 112, E677–E686.

Phillips-Cremins, J.E., Sauria, M.E.G., Sanyal, A., Gerasimova, T.I., Lajoie, B.R., Bell, J.S.K., Ong, C.-T., Hookway, T.A., Guo, C., Sun, Y., et al. (2013). Architectural Protein Subclasses Shape 3D Organization of Genomes during Lineage Commitment. Cell 153, 1281–1295.

Plasschaert, R.N., Vigneau, S., Tempera, I., Gupta, R., Maksimoska, J., Everett, L., Davuluri, R., Mamorstein, R., Lieberman, P.M., Schultz, D., et al. (2014). CTCF binding site sequence differences are associated with unique regulatory and functional trends during embryonic stem cell differentiation. Nucleic Acids Research 42, 774–789.

Rao, S.S.P., Huang, S.-C., Hilaire, B.G.S., Engreitz, J.M., Perez, E.M., Kieffer-Kwon, K.-R., Sanborn, A.L., Johnstone, S.E., Bascom, G.D., Bochkov, I.D., et al. (2017). Cohesin Loss Eliminates All Loop Domains. Cell 171, 305–309.e324.

Rowley, M.J., and Corces, V.G. (2018). Organizational principles of 3D genome architecture. Nature 19, 1–12.

Rudan, M.V., Barrington, C., Henderson, S., Ernst, C., Odom, D.T., Tanay, A., and Hadjur, S. (2015). Comparative Hi-C Reveals that CTCF Underlies Evolution of Chromosomal Domain Architecture. CellReports 10, 1297–1309.

Ruiz-Velasco, M., Kumar, M., Lai, M.C., Bhat, P., Solis-Pinson, A.B., Reyes, A., Kleinsorg, S., Noh, K.-M., Gibson, T.J., and Zaugg, J.B. (2017). CTCF-Mediated Chromatin Loops between Promoter and Gene Body Regulate Alternative Splicing across Individuals. Cell Syst 5, 628– 637.e6.

Saldaña-Meyer, R., González-Buendía, E., Guerrero, G., Narendra, V., Bonasio, R., Recillas-Targa, F., and Reinberg, D. (2014). CTCF regulates the human p53 gene through direct interaction with its natural antisense transcript, Wrap53. Genes & Development 28, 723–734.

Schmidt, D., Schwalie, P.C., Ross-Innes, C.S., Hurtado, A., Brown, G.D., Carroll, J.S., Flicek, P., and Odom, D.T. (2010). A CTCF-independent role for cohesin in tissue-specific transcription. Genome Research 20, 578–588.

Servant, N., Varoquaux, N., Lajoie, B.R., Viara, E., Chen, C.J., Vert, J.-P., Heard, E., Dekker, J., and Barillot, E. (2015). HiC-Pro: an optimized and flexible pipeline for Hi-C data processing. Genome Biology 16, 259.

Sigova, A.A., Abraham, B.J., Ji, X., Molinie, B., Hannett, N.M., Guo, Y.E., Jangi, M., Giallourakis, C.C., Sharp, P.A., and Young, R.A. (2015). Transcription factor trapping by RNA in gene regulatory elements. Science 350, 978–981.

Soto-Reyes, E., and Recillas-Targa, F. (2010). Epigenetic regulation of the human p53 gene promoter by the CTCF transcription factor in transformed cell lines. Oncogene 29, 2217–2227.

Stadhouders, R., Vidal, E., Serra, F., Di Stefano, B., Le Dily, F., Quilez, J., Gómez, A., Collombet, S., Berenguer, C., Cuartero, Y., et al. (2018). Transcription factors orchestrate dynamic interplay between genome topology and gene regulation during cell reprogramming. Nature 50, 238–249.

Weintraub, A.S., Li, C.H., Zamudio, A.V., Sigova, A.A., Hannett, N.M., Day, D.S., Abraham, B.J., Cohen, M.A., Nabet, B., Buckley, D.L., et al. (2017). YY1 Is a Structural Regulator of Enhancer-Promoter Loops. Cell 171, 1–45.

Whyte, W.A., Orlando, D.A., Hnisz, D., Abraham, B.J., Lin, C.Y., Kagey, M.H., Rahl, P.B., Lee, T.I., and Young, R.A. (2013). Master Transcription Factors and Mediator Establish Super-Enhancers at Key Cell Identity Genes. Cell 153, 307–319.

Witcher, M., and Emerson, B.M. (2009). Epigenetic Silencing of the p16INK4a Tumor Suppressor Is Associated with Loss of CTCF Binding and a Chromatin Boundary. Molecular Cell 34, 271–284.

Xie, X., Mikkelsen, T.S., Gnirke, A., Lindblad-Toh, K., Kellis, M., and Lander, E.S. (2007). Systematic discovery of regulatory motifs in conserved regions of the human genome, including thousands of CTCF insulator sites. Proceedings of the National Academy of Sciences 104, 7145– 7150.

Zuin, J., Dixon, J.R., van der Reijden, M.I.J.A., Ye, Z., Kolovos, P., Brouwer, R.W.W., van de Corput, M.P.C., van de Werken, H.J.G., Knoch, T.A., van IJcken, W.F.J., et al. (2014). Cohesin and CTCF differentially affect chromatin architecture and gene expression in human cells. Proc. Natl. Acad. Sci. U.S.a. 111, 996–1001.

