## Supplemental Figures 1-5 for "RNA interactions with CTCF are essential for its proper function"

### SUPPLEMENTARY FIGURES

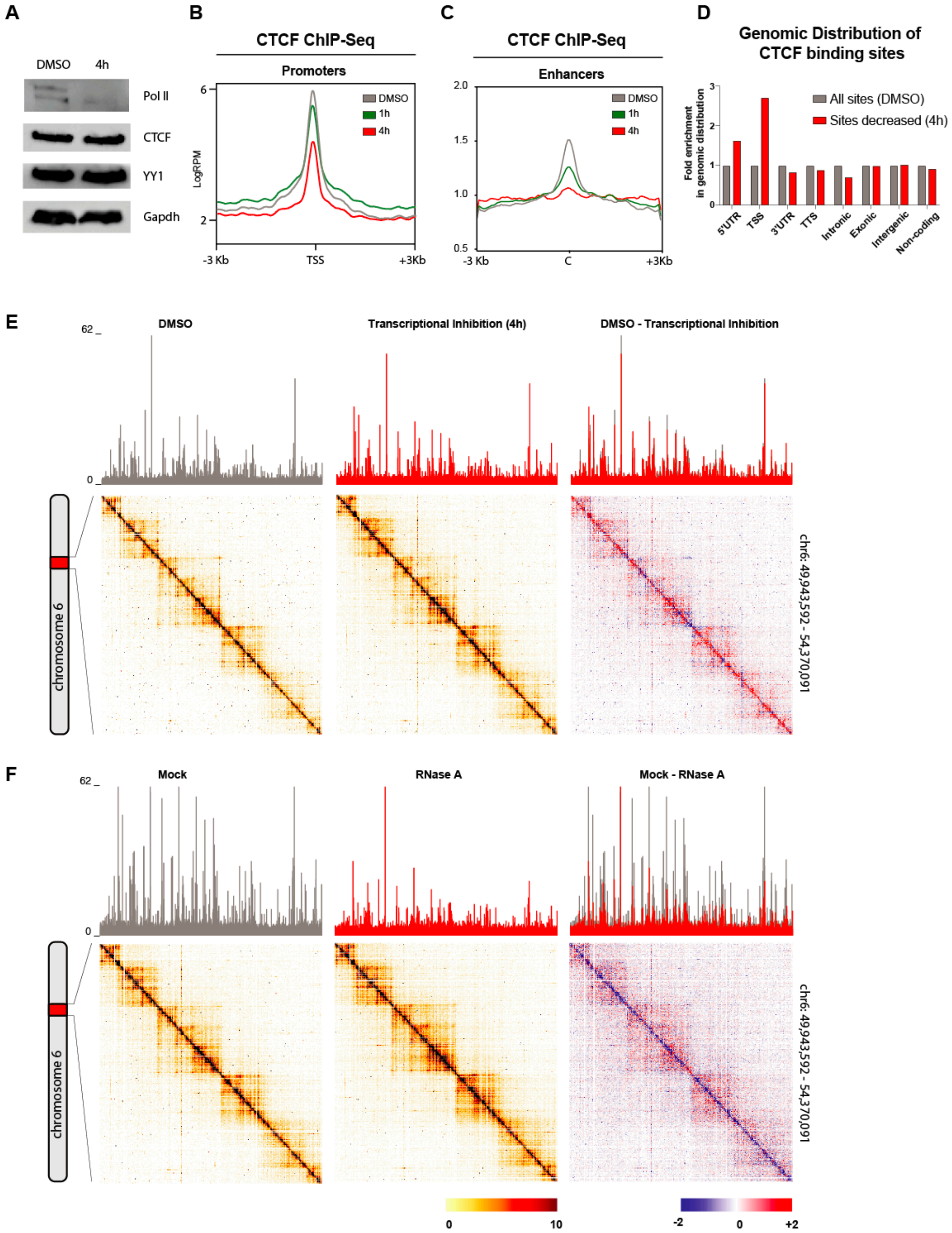


**Figure S1 related to Figure 1. Transcriptional Inhibition disrupts CTCF binding predominantly in TSS.**

mESCs were incubated with a combination of transcriptional inhibitors (Triptolide and DRB) for 4 hours **(A)** Immunoblot for proteins shown in control (DMSO) versus TI (4h). **(B)** Average density profile for CTCF ChIP-Seq centered in the transcriptional start sites of all genes **(C)** or enhancers. **(D)** Bar graph comparing the fold enrichment of the genomic distribution of CTCF binding sites in the control condition against those binding sites significantly decreased TI. **(E-F)** 5C heatmaps depicting the raw interaction frequency between restriction fragments across a 4-Mb region surrounding the HoxA cluster for the TI and RNase A treatments. Data is only normalized to sequencing depth and not binned. Darker colors represent increasing interaction frequency. CTCF ChIP-seq tracks are shown on top. A comparative heatmap is also shown where lost (blue) and gained (red) interactions are illustrated and overlapping ChIP-Seq tracks for control (grey) and treatment (red).

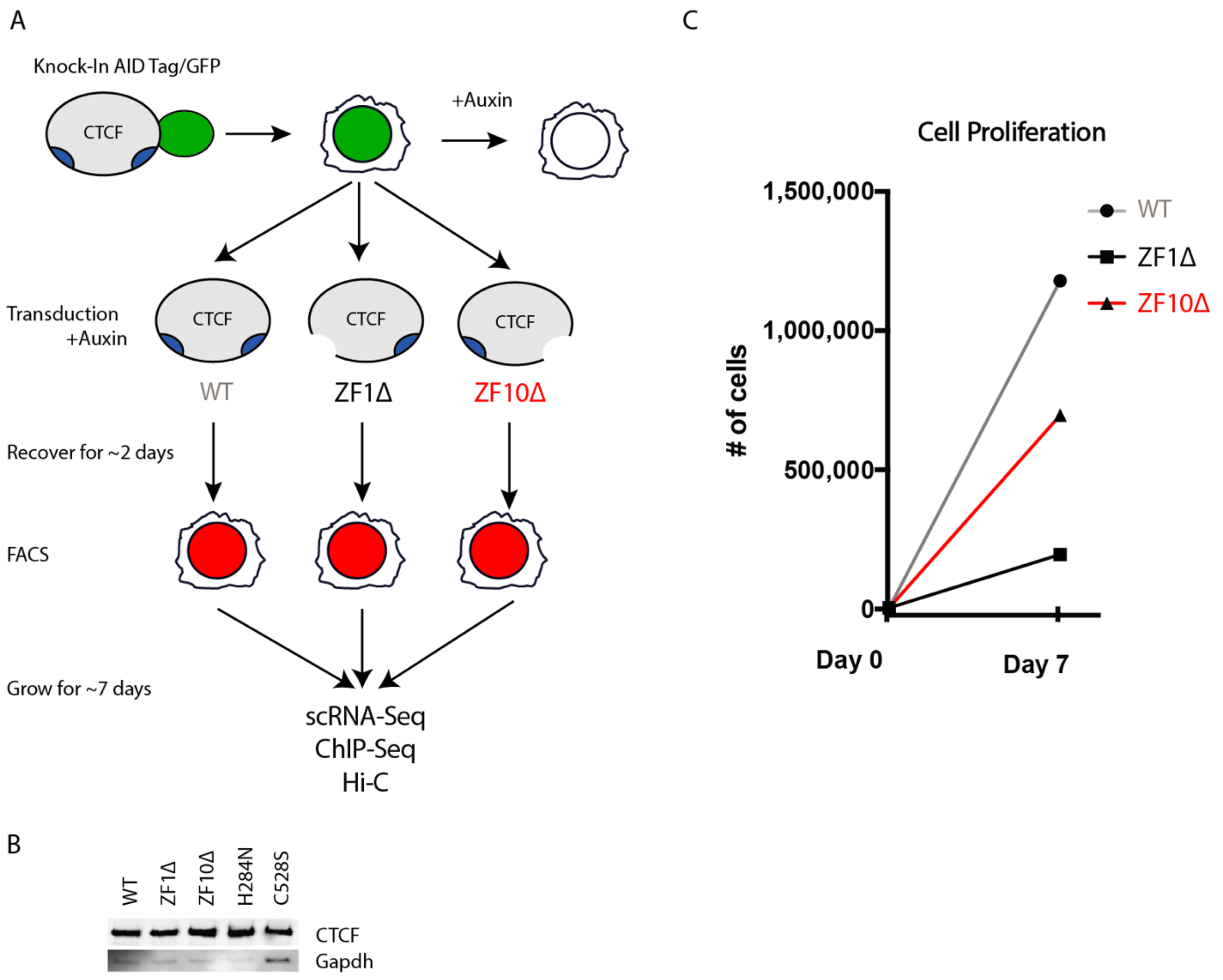


**Figure S2 related to Figure 2. Mutations in ZF1 and ZF10 independently abolish CTCF binding to RNA.**

(A) Schematic representation of the cell lines used. (B) Immunoblot showing CTCF levels in the rescue cell lines indicated. (C) Growth curve showing the difference in proliferation for the different mutants compared to the WT rescue control.

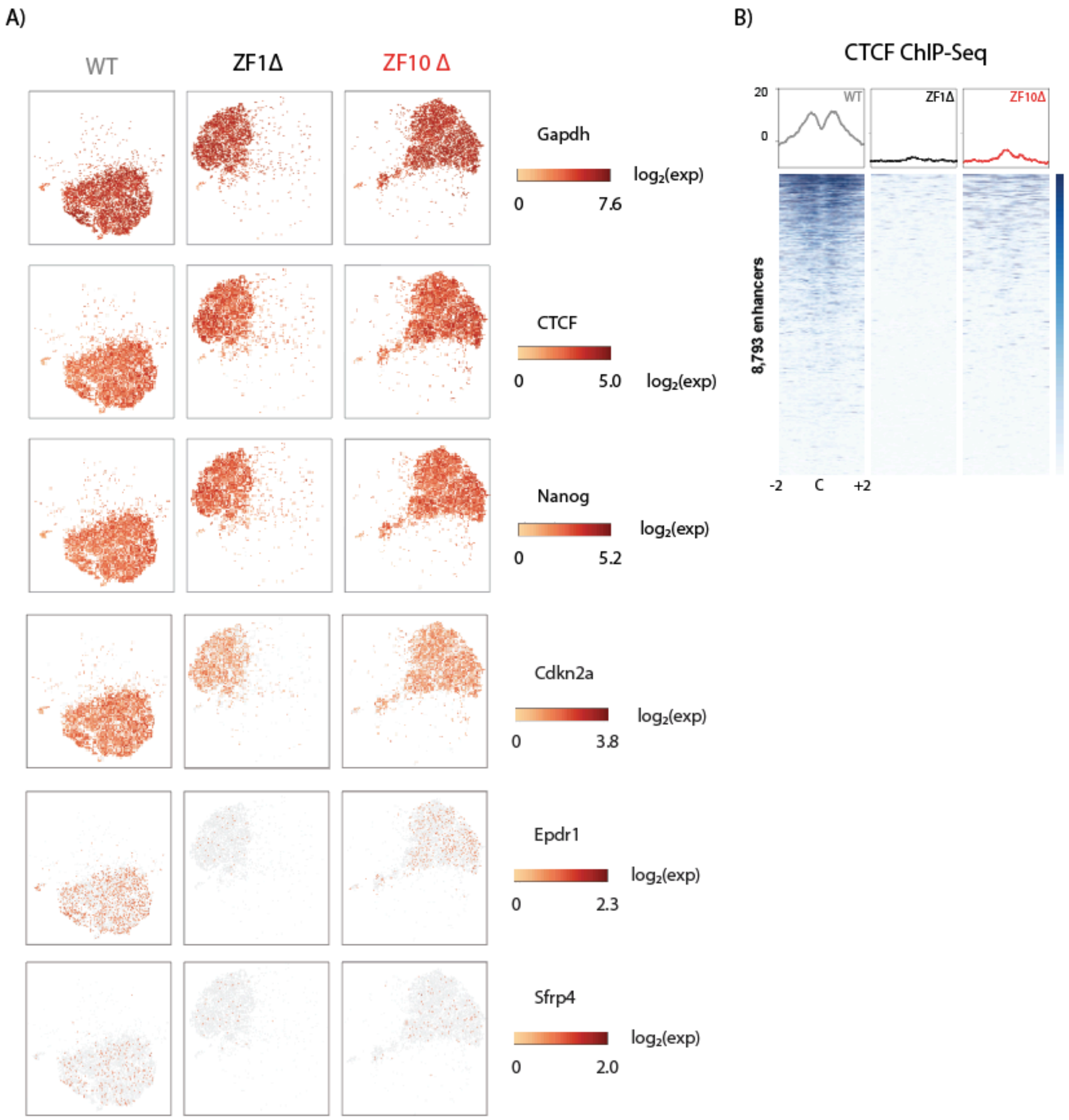


**Figure S3 related to Figure 3. Gene expression defects correlate with CTCF chromatin binding.**

**(A)** Expression the genes shown as measured by single-cell RNA-Seq of rescue cell lines. Each dot represents a single-cell and dots are shaded based on their normalized expression value. **(B)** CTCF ChIP-seq for each cell line indicated. Heatmaps were generated by centering and rank-ordering on enhancers. Corresponding average density profiles are plotted at the top of the heatmaps.

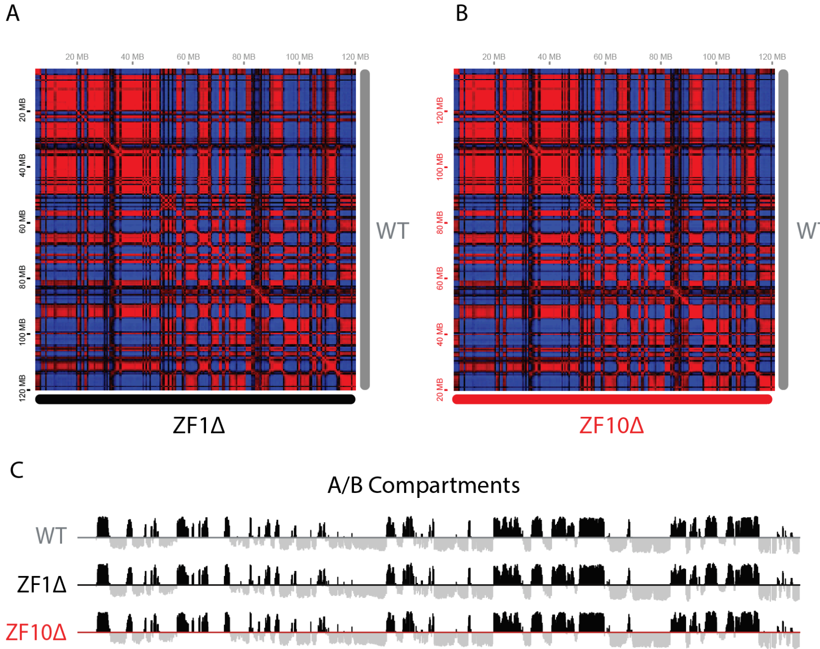


**Figure S4 related to Figure 5. Genomic compartmentalization is unaffected by RBR mutants**
**(A and B)** Representative Pearson's correlation maps ([A], ZF1 vs WT; [B], ZF10Δ vs WT) of chromosome 12 showing the plaid pattern fine-scale compartmentalization is unaffected. **(C)** Comparison of distributions of cis-Eigenvector 1 values across chromosome 12 for all conditions.

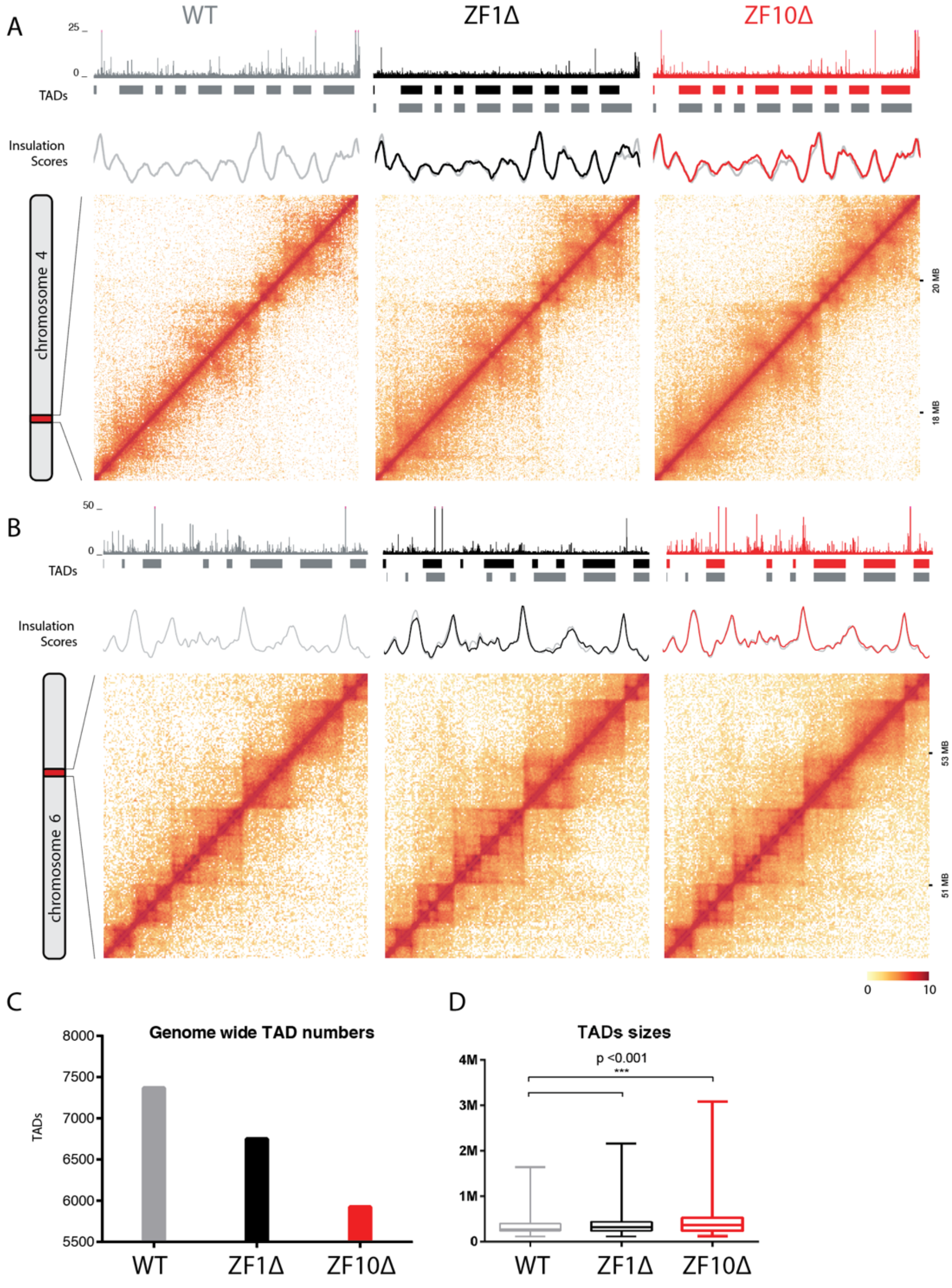


**Figure S5 related to Figure 5. RBR mutants correlates with a global weakening of boundaries.**
**(A-B)** Hi-C contact maps at 20 kb resolution for 2 representative regions. From top to bottom, ChIP-Seq tracks for CTCF are shown, then TADs and finally insulation scores graphed along the region. **(C)** Number of TADs identified through IS with a 40kb resolution and a 102 kb window size. The cut-off used was adjusted to the WT sample and used for all experimental conditions. **(D)** TADs sizes vary between WT and ZF1 and ZF10 mutants. Box whiskers plot showing the median and the 2.5-97.5 percentile of the TAD size distribution. Significant differences are detected between WT and ZF1 and ZF10 mutants TAD sizes by Wilcoxon Ranked Test.
